# Leveraging a self-cleaving peptide for tailored control in proximity labeling proteomics

**DOI:** 10.1101/2023.11.03.565112

**Authors:** Louis Delhaye, George D. Moschonas, Daria Fijalkowska, Annick Verhee, Delphine De Sutter, Tessa Van de Steene, Margaux De Meyer, Laura Van Moortel, Karolien De Bosscher, Thomas Jacobs, Sven Eyckerman

## Abstract

Protein-protein interactions play an important biological role in every aspect of cellular homeostasis and functioning. Proximity labeling mass spectrometry-based proteomics overcomes challenges typically associated with other methods, and has quickly become the current state-of-the-art in the field. Nevertheless, tight control of proximity labeling enzymatic activity and expression levels is crucial to accurately identify protein interactors. Here, we leverage a T2A self-cleaving peptide and a non-cleaving mutant to accommodate the protein-of-interest in the experimental and control TurboID setup. To allow easy and streamlined plasmid assembly, we built a Golden Gate modular cloning system to generate plasmids for transient expression and stable integration. To highlight our T2A Split-link design, we applied it to identify protein interactions of the glucocorticoid receptor and SARS-CoV-2 nucleocapsid and NSP7 proteins by TurboID proximity labeling. Our results demonstrate that our T2A split-link provides an opportune control that builds upon previously established control requirements in the field.

**Motivation:** In proximity labeling proteomics protein-protein interactions are identified by *in vivo* biotinylation. However, the current lack of a universally applicable negative control for differential analysis affects accurate mapping of the interactome. To bridge this gap, we conceptualized a system based on the T2A self-cleaving peptide to match expression levels between control and bait protein setups while using the same bait protein. In addition, we implemented a versatile modular cloning system to build mammalian expression vectors for, but not limited to, proximity labeling assays.

## INTRODUCTION

Proximity labeling proteomics has revolutionized protein-protein interaction (PPI) discovery. Enzymes such as BirA^1^, HRP^2^, APX^3^, Ubc12^4^, and PafA^5^ have all been engineered to allow promiscuous labeling of proteins in live cells. These enzymes can be genetically attached to a bait protein to covalently label proteins in close proximity. Especially BirA (BioID) and APX (APEX2^6^) have been widely used. Whereas the original BioID utilizes a R118G mutant version of the *E. coli* biotin ligase BirA, termed BirA*, more recent derivatives such as TurboID^7^ have been engineered using directed evolution. TurboID allows for shorter labeling times (10 min to a few hours) compared to the original BioID (15-18 hours^1^) making it preferable for interaction dynamics and temporal control. Both BioID and APEX2 catalyze the covalent attachment of biotin handles to certain amino acid side chains upon supplementation of a biotin substrate. After lysis, biotinylated proteins are captured by streptavidin enrichment. The *in vivo* labeling permits stringent lysis and washing conditions that effectively nullify the loss of interactors due to post-lysis dissociations caused by lowering the concentration of protein partners upon lysis. In addition, these harsh conditions prevent post-lysis associations due to aspecific protein binding to the affinity resin and associated reagents, as well as to the formation of non-physiological PPIs caused by loss of subcellular compartimentalization^8,9^. Therefore, these harsh conditions reduce the number of false positives compared to antibody-based affinity purification approaches. In contrast, due to the promiscuous nature of the labeling enzyme, non-interacting bystander proteins can also be labeled and act as false positives. To limit the extent of false positives identified this way, a quantitative proteomics approach with a suitable negative control has to be employed. Such controls should contain a comparable labeling activity, and be (partially) located in the same subcellular location to match the proteome that can be labeled. Most notably, to match labeling activity quantitatively, expression levels of the labeling enzyme need to be equivalent across all conditions. Typical used negative controls include irrelevant bait proteins, such as GFP, tagged with the labeling enzyme, or a free untagged labeling enzyme. Although expression levels can be matched by inducible dose-dependent promoters, different properties of these irrelevant proteins, such as the tendency to aggregate, can severely impact the identified interactome. In addition, the protein of interest might induce specific changes in the global proteome, e.g. transcription factors, that may or may not affect the interactome. As the aforementioned negative controls setups are unlikely to induce these global proteomic changes or to the same extent, significantly enriched interactors might simply reflect differences in global proteomes between both setups rather than specific interactomes.

To overcome these limitations, we extended on our previous design to use a T2A self-cleaving peptide for proximity labeling of endogenous bait proteins^10^. Inclusion of the T2A peptide in a coding sequence causes ribosomal skipping of the G-P peptide bond at the C-terminus of the peptide, effectively generating a translational polycistron in which the proteins up- and downstream of the T2A peptide are physically separated, yet translated at equimolar amounts^11,12^. We reasoned that engineering a control cell line containing an inactivated T2A (MUTT2A) sequence would provide the opportunity to use the same bait protein for the experimental and control setup (Fig. 1). This strategy overcomes the challenges described before. To permit customizable and rapid plasmid assembly of proximity labeling constructs for mammalian expression, we built a modular cloning system based on Golden Gate assembly to generate plasmids for lentiviral and transposon-based genomic integration of transgenes in mammalian cell lines.

**Fig. 1.**
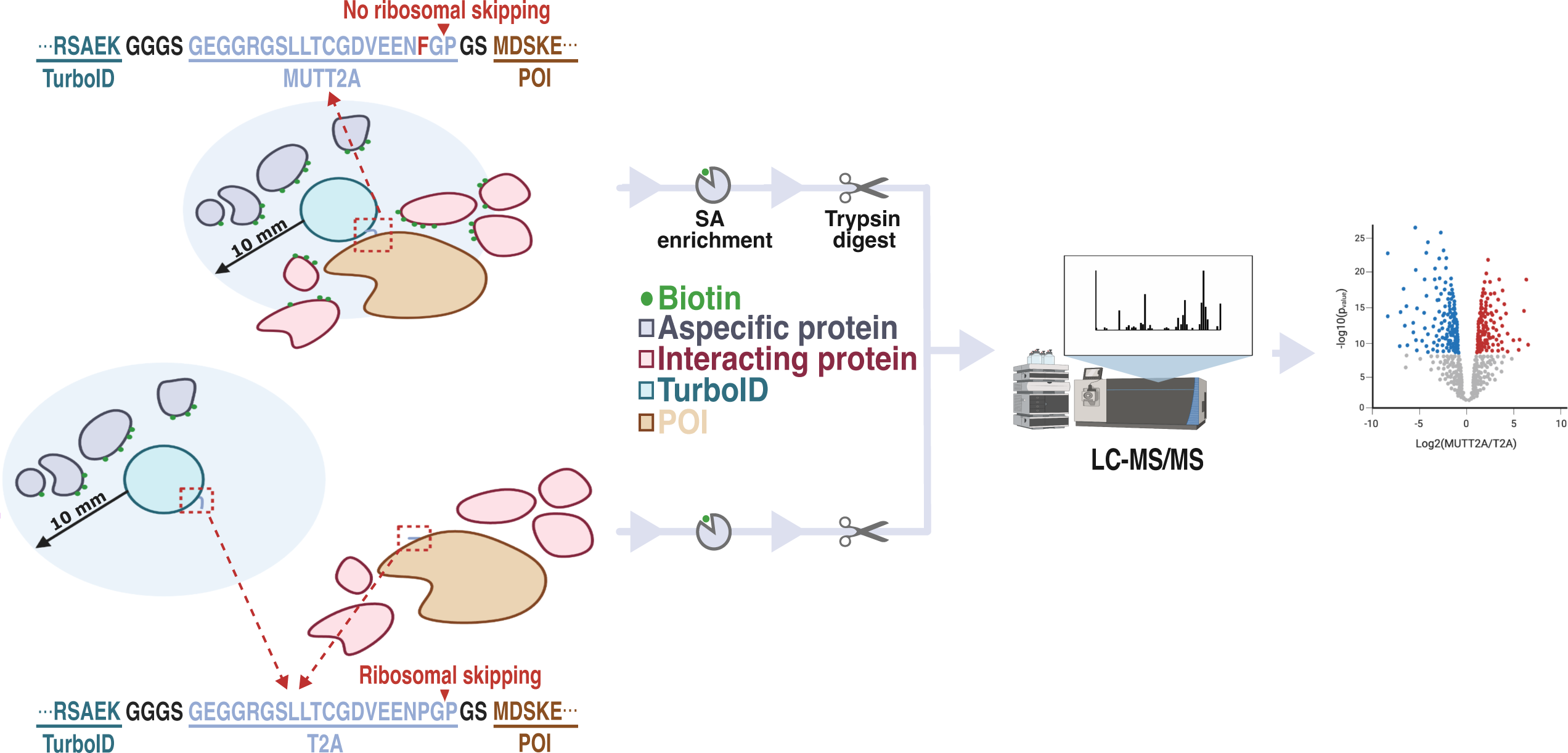
Workflow of a T2A split-link design. Engineered T2A and MUTT2A cell lines express TurboID (cyan) and the POI (brown) either physically separated or as a fusion protein, respectively. Proximal proteins in either setup are enriched by streptavidin pulldown, and quantified and identified by LC-MS/MS. Differential analysis between both setups highlights POI interaction partners as proteins enriched in the MUTT2A setup. POI, protein-of-interest; SA, streptavidin; LC-MS/MS, liquid chromatography with tandem mass spectrometry.

## RESULTS

### Establishing a Golden Gate modular assembly platform for mammalian expression constructs

Golden GateWAY and GreenGate utilize a sequential cloning approach of Golden Gate and Multisite Gateway^TM^ cloning to rapidly generate complex expression vectors^13,14^. However, the system does not allow for mammalian expression. Therefore, as a starting point, we adjusted the Golden GateWAY (GGW) and GreenGate platforms to be compatible with mammalian expression. As is common in modular cloning platforms, GGW contains a hierarchy of three levels. First six modules (Level-0) are cloned in a one-pot BsaI-based Golden Gate assembly mix into transcriptional units (TU; Level-1). Modules are flanked by BsaI recognition sites that generate unique overhangs, termed A to G, depending on the component that the module includes. The unique overhangs for each position allow unidirectional assembly of the modules. We adopted these overhangs, but slightly changed the content of certain positions. Rather than including a positive selection cassette, as in Golden GateWAY and GreenGate^13,14^, we opted for repurposing the FG position to comprise a transcriptional terminator sequence (e.g. pA signals) or a post-transcriptional element (e.g. WPRE sequence). As a result, DE and EF were adopted to respectively include a linker sequence and a C-terminal tag instead of a C-terminal tag and terminator sequence as is the case in GreenGate.

Level-1 plasmids contain attL and attR sites that are situated up- and downstream of the TU to allow LR Multisite Gateway^TM^ reactions for assembly of multiple TUs in the final expression construct (Level-2). We repurposed pEN-L4-AG-R1^15^, pEN-L1-AG-L2^15^, and pEN-R2-AG-L3^16^ as Level-1 destination vectors. In addition, we generated pEN-L1-AG-L3 to accommodate combining two (L4-R1 & L1-L3) or three (L4-R1 & L1-L2 & R2-L3) different TUs within a R4-R3 Level-2 destination vector. As lentiviral transduction still presents one of the most straightforward and robust methods to stably integrate a transgene into a mammalian cell, we built a Level-2 destination vector, termed pLV-GGW-DEST, by inserting the R4-R3 cassette from pMG426^17^ between the necessary regulatory lentiviral sequences and the 3’ long terminal repeat (LTR) of a 3^rd^ generation SIN lentiviral vector. Using our mammalian expression-compatible Golden GateWAY platform, as a proof-of-concept, we assembled a lentiviral vector that expresses monomeric superfolder (msf)GFP and nuclear mCherry fluorescent reporters, and allows selection by puromycin after lentiviral transduction (Fig. S1a). Level-0 modules were generated from in-house constructs or by DNA synthesis. Lentiviral particles were produced and subsequently used to transduce human SK-N-BE(2)-C and SHEP neuroblastoma cell lines. Both cell lines conferred puromycin resistance, and the subcellular localization of both fluorescent signals was consistent with their localization signals (Fig. S1b). This demonstrates our cloning platform is capable of generating fully custom lentiviral transfer vectors for transgene integration in mammalian cells.

Because lentiviral vectors are recombination-prone^18^ and Gateway^TM^ reagents are relatively expensive for high-throughput cloning efforts, we decided to swap the LR recombinase step for a second BsmBI-based Golden Gate assembly step (Fig. 2). To do so, we generated a high copy number plasmid with kanamycin resistance that contained BbsI restriction sites to clone between. We removed two BsmBI restriction sites in the bacterial backbone of the plasmid, and inserted a W-A-G-X, X-A-G-Y, Y-A-G-Z, or a X-A-G-Z cassette between the BbsI restriction sites. Similar to the BsaI overhangs, W, X, Y, and Z are unique overhangs that are generated by restriction digest with BsmBI allowing Golden Gate assembly of higher order plasmids. These overhangs were chosen based on Potapov et al. ^19^ for having a high ligation fidelity with T4 DNA ligase and no observable ligation at possible DNA base pair mismatches with any of the other overhangs. Similar as pLV-GGW-DEST, we inserted a W-Z cassette between the LTRs of the same lentiviral vector to construct pLV-W-Z. In addition, we inserted a W-Z cassette between the 5’ and 3’ inverted terminal repeats (ITR) of AAT-PB-CG2APtk^20^ to allow Golden Gate assembly of piggyBac transposable vectors for genomic integration in mammalian cells. All A-G and W-Z cassettes encode a chloramphenicol resistance gene (CmR; *cat* gene) and *ccdB* toxin gene expressed by a strong, constitutive *lac*UV5 bacterial promoter to provide positive or negative selection, respectively. Moreover, Level-0 and Level-2 vector backbones confer carbenicillin or ampicillin resistance (AmpR) through expression of a *bla* gene, while Level-1 vector backbones confer kanamycin resistance (KanaR) through expression of an *aph* gene to prevent selection of transformants containing lower-ordered plasmids within the hierarchy.

**Fig. 2.**
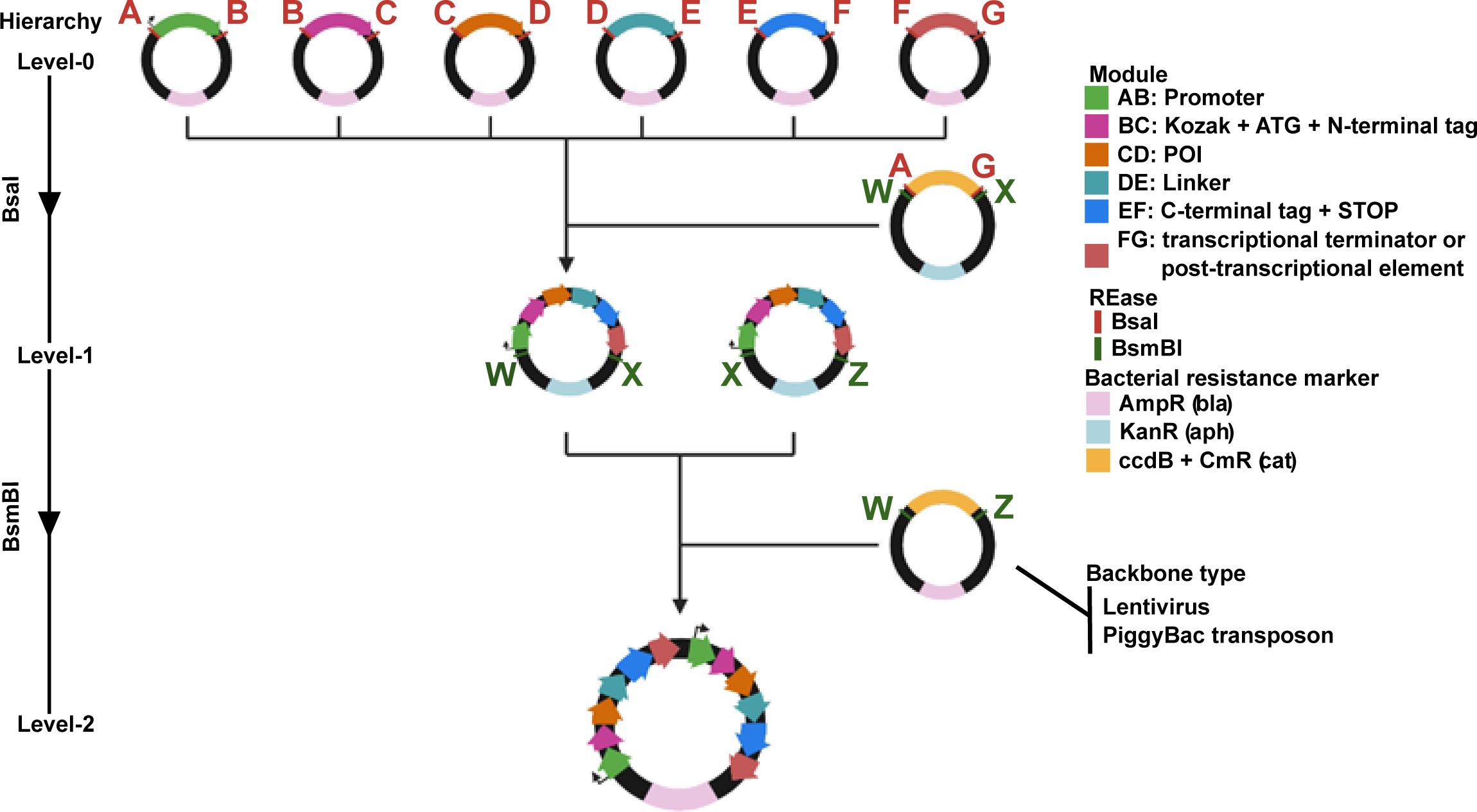
A mammalian Golden Gate assembly system. Plasmids are arranged in hierarchy depending on their content. Level-0 are basic parts flanked by inward-cutting BsaI restriction sites. BsaI-based Golden Gate cloning of six Level-0 plasmid (with each position present) and a A-G backbone causes unidirectional assembly of each of these parts in the backbone plasmid. These Level-1 plasmids express transcriptional units, and can similarly be assembled in a higher-order Level-2 plasmid containing two Level-1 transcriptional units by BsmBI-based Golden Gate assembly.

### T2A split-link design identifies the glucocorticoid receptor interactome

As a first experiment, we performed a T2A split/link proximity labeling screen with TurboID for the glucocorticoid receptor (GR, gene symbol: *NR3C1*). GR is a nuclear receptor that is sequestered in the cytoplasm but relocates to the nucleus upon glucocorticoid (GC) binding. There it acts as a transcription factor to regulate target gene expression. Using our Golden Gate assembly platform, we generated TurboID-T2A/MUTT2A-GR expression vectors with a dose-dependent doxycycline-responsive (TRE) promoter (Fig. S2a). These constructs were combined with a second TU expressing a blasticidine resistance gene within a piggyBac transposon backbone. We tagged GR N-terminally with V5-TurboID-T2A/MUTT2A, as tagging the C-terminus might impair ligand binding and thus proper GR function^21^. Human lung epithelial A549 cells constitutively expressing a tetracycline-inducible transactivator were co-transfected with either T2A or MUTT2A TurboID-GR constructs, and piggyBac transposase (PBase) to generate stable A549-TurboID-(MUT)T2A-GR cell populations. As the number of transposition events in the T2A and MUTT2A setup might be different between both cell lines, we matched expression levels of both transgenes by assessing a range of doxycycline to equalize the amount of TurboID and GR present in the cell (Fig. S2b). Indeed, we found 20 ng mL^−1^ and 150 ng mL^−1^ doxycycline to express an equal amount of TurboID and biotinylation in the T2A and MUTT2A setups, respectively. We observed no ribosomal skipping in the MUTT2A cell line, whilst we observed no full length TurboID-GR fusion proteins in the T2A cell line (Fig. S2b). At these near-physiological expression levels, we also observed a similar upregulation of well-known anti-inflammatory GR target genes (*TSC22D3* and *DUSP1*)^22^ upon dexamethasone supplementation, a potent GR agonist, indicating both cell lines still trigger activation of the same downstream targets and show a comparable GR-dependent transcriptional transactivation (Fig. S2c).

After validating the cell lines, we performed proximity labeling after supplementation of dexamethasone at the doxycycline concentrations we determined earlier. An outlying replicate was removed based on principal component analysis (PCA, Fig. S2d). Differential analysis showed 292 proteins to be significantly enriched in the MUTT2A samples at a 5% FDR (Fig. 3a, Table S1). We retrieved well-known GR coactivators NCOA2, NCOA3, and NCOA6, as well as components of chromatin remodeling complexes such as SWI/SNF. iBAQ intensities of endogenously biotinylated proteins were comparable between both setups, demonstrating similar enrichment efficiencies (Fig. S2e). Similarly, iBAQ intensities of housekeeping genes (PPIA, TUBB, YWHAZ, GAPDH, VIM) were either comparable or enriched in the T2A setup, demonstrating the aspecific background was consistent between both setups (Fig. S2f). In contrast, we identified a significant enrichment of TurboID in the MUTT2A samples (Fig. 3a), something we previously also saw in our TP53-T2A/MUTT2A-BioID study^10^. We performed gene set enrichment analysis (GSEA) preranked by LOG2FC with the BioID data of Lempiainen et al.^23^ and Dendoncker et al. ^24^ as gene sets (Fig. 3b). Interactors identified in both studies were significantly enriched in our data set (Table S1, adj. P-value = 2.07 x 10^−9^ and 2.07 x 10^−9^, respectively), highlighting our results are consistent with previous proximity biotinylation interactome studies performed for GR. Moreover, we queried the interactors to the C2 (v2023.1) and C5 (v2023.1) collection of the Molecular Signatures Database (MSigDB). Overrepresented pathways in our data were in line with previously published results, such as crosstalk with PPARα^25,26^ and AR^23^, or GR’s known role in circadian biology^27–29^ (Fig. 3c). Top overrepresented ontology terms included the mediator complex and RNAPII preinitiation complex assembly (Fig. 3c), terms expected to be overrepresented for an active transcription factor.

**Fig. 3.**
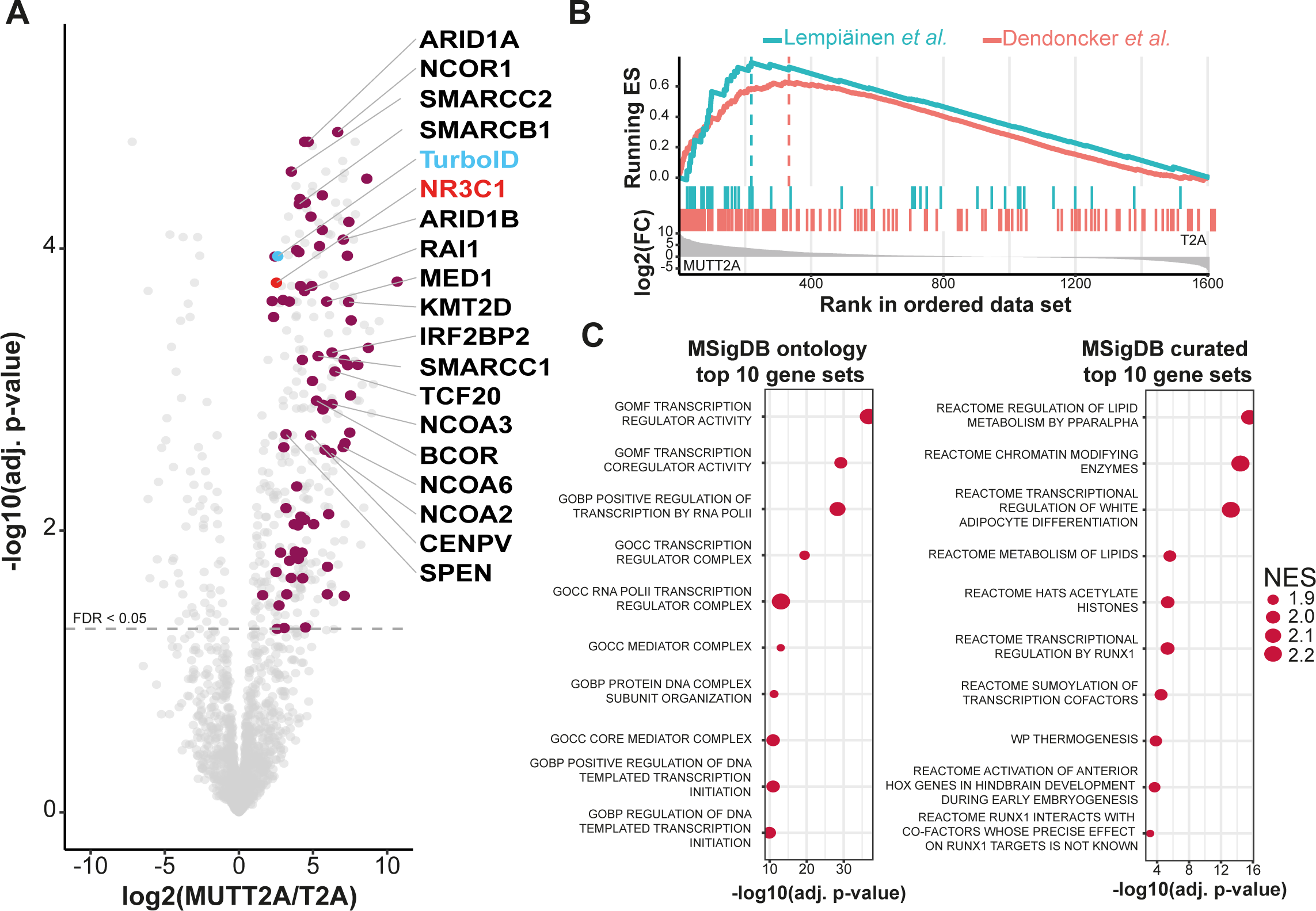
Proximity labeling of the glucocorticoid receptor with a T2A split-link design. (a) differential analysis of TurboID-T2A-GR and TurboID-MUTT2A-GR. Bait is shown in red, TurboID in blue. Significant (adj. P-Value < 0.05) proteins that are also found in BioGRID are shown in crimson. Highlighted proteins are proteins also identified in other proximity labeling studies. (b) Pre-ranked gene set enrichment analysis (GSEA) of other proximity labeling studies within our data set. Maximal running enrichment score for each study is highlighted by a dotted line. (c) Pre-ranked GSEA showing enriched pathways in the c2 (curated gene sets) and c5 (ontology) collections of the Molecular Signature Database (MSigDB). Top 10 pathways enriched in the MUTT2A setup are shown. NES, normalized enrichment score.

### SARS-CoV-2 nucleocapsid protein interactors are enriched for stress granule components

After applying our T2A split-link design to GR, we applied the same setup to SARS-CoV-2 nucleocapsid protein (NCAP). NCAP binds and shields the viral genome of SARS-CoV-2, and has been shown to counter antiviral host responses upon infection^30^. Moreover, multiple studies^31–33^ and preprints^34,35^ have provided an NCAP interactome by proximity labeling, allowing us to compare with our T2A split-link design. Using our Golden Gate assembly platform, we generated both N- and C-terminally (MUT)T2A-TurboID-tagged NCAP piggyBac-compatible constructs (Fig. S3a) and stably integrated them in A549 cells that expressed the tetracycline-inducible transactivation machinery. Similar as for GR, all constructs were under the control of a doxycycline-inducible promoter and expression levels were assessed by immunoblotting over a range of doxycycline. We observed that C-terminal tagged T2A and MUTT2A NCAP setups did not contain comparable amounts of TurboID which was also reflected in differing amounts of biotinylation between the T2A and MUTT2A setups (Fig. S3b). Therefore, we proceeded with the N-terminally tagged cell lines for further experiments (Fig. S3c). We found 25 ng mL^−1^ doxycycline to have equal amounts of biotinylation and used this concentration for all subsequent experiments. Interestingly, at this concentration TurboID amounts seemed slightly lower in the MUTT2A setup. However, we observed additional TurboID staining at a lower molecular weight, consistent with previously reported alternative N-terminal processing of NCAP by cellular proteases^36^. The combined pool of TurboID is very likely similar between both setups and corroborates the equal amounts of biotinylation.

Next, we performed proximity labeling with the conditions as described above. An outlying replicate was removed based on PCA analysis (Fig. S3d). Differential analysis between the T2A and MUTT2A setups resulted in 51 significantly enriched proteins at a 5% FDR (Fig. 4a, Table S2). With the exception of ACACA, iBAQ intensities of endogenously biotinylated proteins (Fig. S3e) and house-keeping genes (Fig. S3f) were comparable between both setups, demonstrating similar enrichment efficiencies and a similar aspecific background. Similar as with GR, we saw a significant enrichment of TurboID in the MUTT2A compared to the T2A condition (Fig. 4a). Using GSEA, we compared our results with other proximity labeling studies for NCAP and found our NCAP interactors to be significantly enriched (Table S2, adj. P-value = 2.4 x 10^−9^, 2.7 x 10^−8^, 8.6 x 10^−8^, 2.8 x 10^−5^, and 0.002 for Laurent *et al*.^35^, Liu *et al*.^31^, Samavarchi-Tehrani *et al*.^34^, May *et al.*^33^ and Zhang *et al*.^37^, respectively) in these studies (Fig. 4b). Core stress granule (SG) components were among the most significantly enriched proteins. Moreover, we also identified GSK3B, a subunit of the GSK3 kinase that is known to phosphorylate NCAP^38,39^. These results are consistent with previous PPI studies and NCAP’s function to attenuate the antiviral innate immune response by preventing SG formation and RIG-I-like receptor signaling activation^40,41^.

**Fig. 4.**
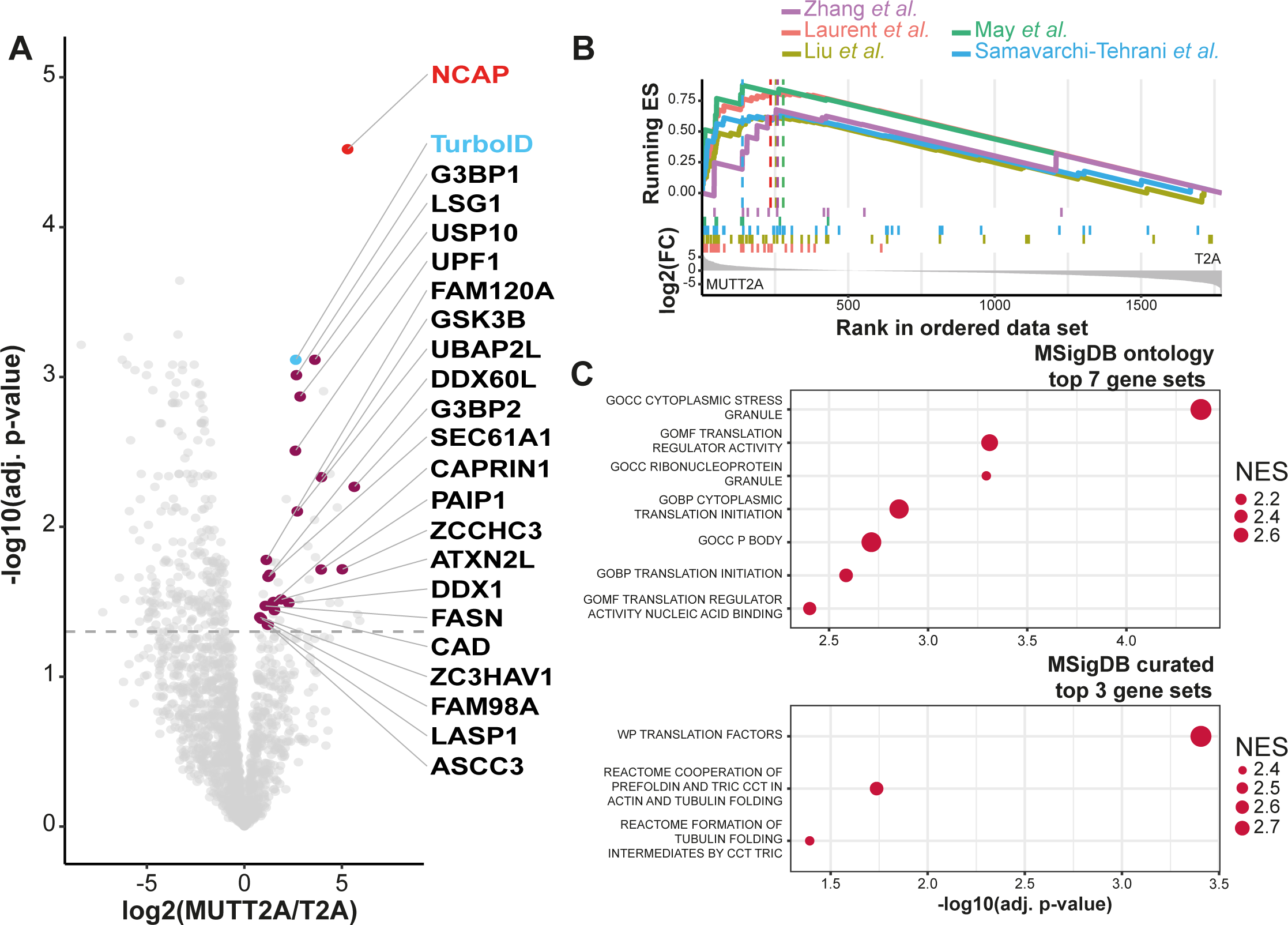
Proximity labeling of SARS-CoV-2 NCAP with a T2A split-link design. (a) differential analysis of TurboID-T2A-NCAP and TurboID-MUTT2A-NCAP. Bait is shown in red, TurboID in blue. Significant (adj. P-Value < 0.05) proteins that are also found in BioGRID are shown in crimson. (b) Pre-ranked gene set enrichment analysis (GSEA) of other proximity labeling studies within our data set. Maximal running enrichment score for each study is highlighted by a dotted line. (c) Pre-ranked GSEA showing enriched pathways in the c2 (curated gene sets) and c5 (ontology) collections of the Molecular Signature Database (MSigDB). Top pathways enriched in the MUTT2A setup are shown. NES, normalized enrichment score.

### T2A split-link identifies SARS-CoV-2 NSP7 interactors

As a final model, we applied our T2A split-link approach on SARS-CoV-2 non-structural protein 7 (NSP7). NSP7 is a 83 amino acid polypeptide that plays an integral role in the transcription of the viral genome. Together with NSP8 and NSP12, it forms the RNA-dependent RNA polymerase (RdRp) supercomplex. Similarly to NCAP, we integrated and assessed N- and C-terminal NSP7- TurboID fusions with a T2A or MUTT2A setup in A549-tetracycline inducible transactivator cells (Fig. S4a). For N-terminal fusion proteins, we observed the expected bands (Fig. S4b), while C-terminal fusions expressed poorly with barely any detectable expression in the MUTT2A setup (Fig. S4c). Biotinylation patterns demonstrated a similar trend (Fig. S4b,c). We did not observe a doxycycline-dependent increase for the N-terminal fusion in either protein expression or biotinylation patterns, with an overall higher expression in the T2A setup (Fig. S4c). This suggests that the operator sites of the TRE promoter were saturated even at the lowest doxycycline concentration, consistent with a low amount of integrations. Therefore, both setups were induced with 25 ng mL^−1^ doxycycline for any downstream experiments.

After removal of outlying replicates by PCA (Fig. S4d), differential analysis found 23 significantly enriched proteins at a 5% FDR (Fig. 5a, Table S3). Both endogenously biotinylated proteins (Fig. S4e) and house-keeping proteins (Fig. S4f) were comparable between both setups, demonstrating similar amounts of enrichment and a common aspecific background in both setups. Except for the data set of Samavarchi-Tehrani et al. ^42^, the small amount of enriched proteins, and, therefore, small overlap did not allow us to perform a GSEA comparison with other proximity labeling studies found in BioGRID. Nonetheless, there was a small yet significant (Table S3, adj. P-value = 0.005) overlap between Samavarchi-Tehrani *et al.* ^42^ and proteins enriched in the MUTT2A setup of our data set (Fig. 5b). Consistent with literature^43–45^, enriched MSigDB c2 and c5 gene sets included sets involved in mitochondrial metabolism and several metabolic processes, respectively (Fig. 5c).

**Fig. 5.**
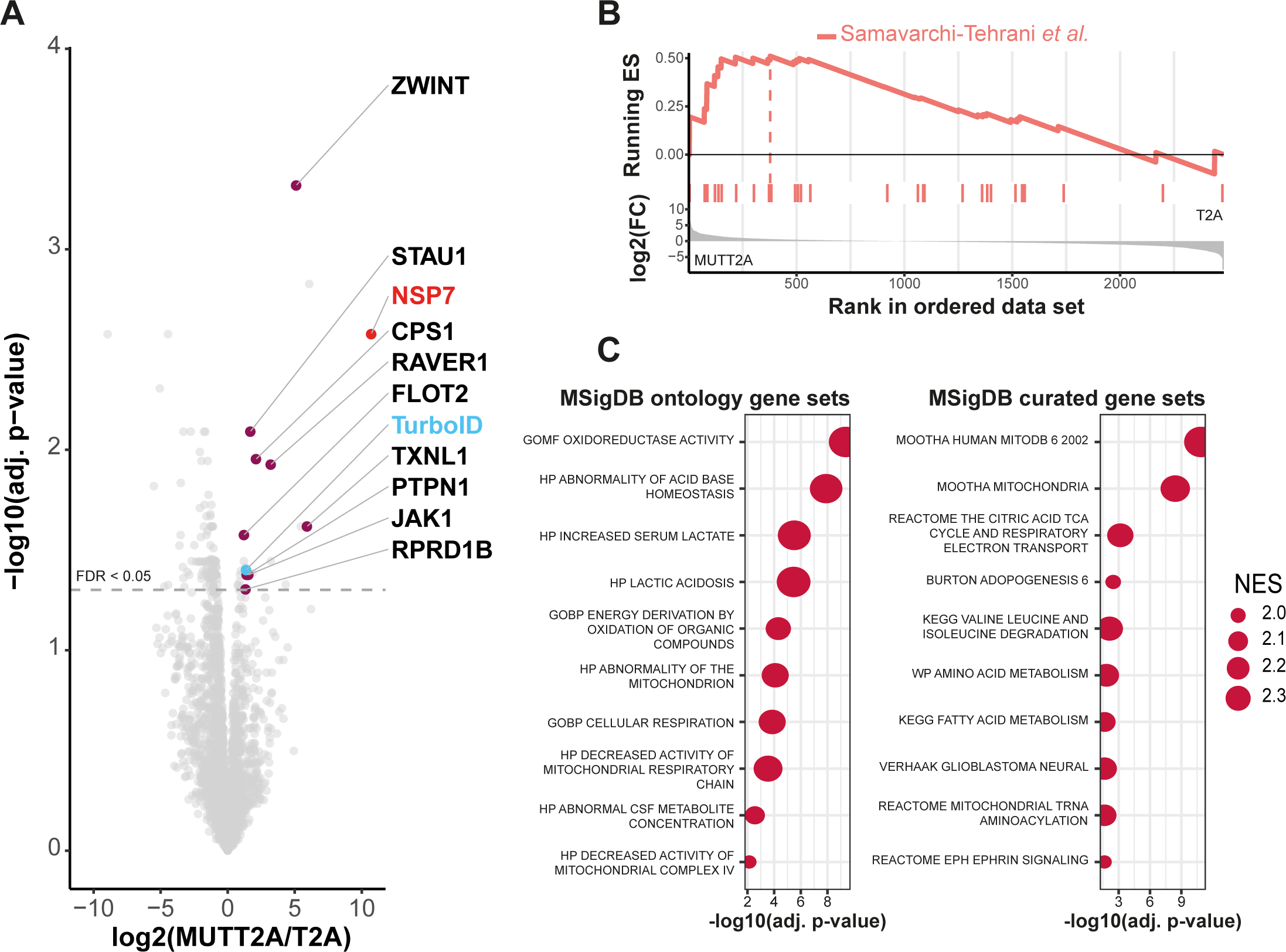
Proximity labeling of SARS-CoV-2 NSP7 with a T2A split-link design. (a) differential analysis of TurboID-T2A-NSP7 and TurboID-MUTT2A-NSP7. Bait is shown in red, TurboID in blue. Significant (adj. P-Value < 0.05) proteins that are also found in BioGRID are shown in crimson. (b) Pre-ranked gene set enrichment analysis (GSEA) of other proximity labeling studies within our data set. Maximal running enrichment score for each study is highlighted by a dotted line. (c) Pre-ranked GSEA showing enriched pathways in the c2 (curated gene sets) and c5 (ontology) collections of the Molecular Signature Database (MSigDB). Top 10 pathways enriched in the MUTT2A setup are shown. NES, normalized enrichment score.

### Subcellular localization of TurboID and the POI determine differences in T2A differential proteins

We noticed that our NCAP and NSP7 data sets overall contained more differential proteins in the T2A setup compared to our GR data set (Fig. S5a). Moreover, overlapping significant proteins enriched in each T2A setup demonstrates the T2A enriched proteomes are more similar between NCAP and NSP7 (Fig. S5b). In contrast, the overlap with the GR experiment was very limited. We wondered whether differences in subcellular localization of the TurboID biotin ligase in the T2A compared to the MUTT2A cell line can explain T2A differences between differential analyses for different POIs. Therefore, we mapped the subcellular localization of all significant proteins based on the immunofluorescence data of the Human Protein Atlas. Indeed, most of the significantly differential proteins in the T2A setup of our NCAP and NSP7 data sets were nuclear (Fig. S5c). In contrast, in our GR experiment most of the significant proteins in the T2A setup were cytosolic. Interestingly, for the MUTT2A side we generally observed the opposite, with most of the significant proteins being nuclear or cytosolic for GR and NCAP, respectively. For NSP7, the amount of significantly differential proteins in the MUTT2A setup is too low to make definite conclusions. We hypothesized that free TurboID (as in the T2A setup) would be present in both compartments, while in the MUTT2A setup, the subcellular localization would be based on the POI. Consistent with these observations, we and others have previously shown that the BirA* biotin ligase can passively diffuse into the nucleus^10,46^, which our data here would suggest is also the case for free TurboID. Indeed, NCAP and NSP7 reside mostly in the cytosol^38,40,47^, while dexamethasone-activated GR would be solely nuclear. To substantiate our claims, we performed immunofluorescence experiments on T2A/MUTT2A cell lines for GR and NCAP (Fig. S5d). In either T2A cell line, we saw a clear distribution of free TurboID over the entirety of the cell, while for MUTT2A the subcellular localization of TurboID was dependent on the POI. Dexamethasone-activated GR restricted TurboID and, therefore, biotinylation to the nucleus, while the NCAP fusion protein and biotinylation activity were only observed in the cytosol. Taken together, we argue that the observed T2A skew in our data can be explained by differential TurboID localization between the T2A and MUTT2A setup. In addition, our GR data did not demonstrate a similar skew, which is in line with an activated transcription factor residing in the nucleus.

### MUTT2A/T2A differential analysis recapitulates a stable BioID background

To assess whether a comparable background in our T2A/MUTT2A setup compared to other BioID experiments can be retrieved, we looked at the iBAQ intensities of TOP1, PARP1, PKM, PRKDC, FLNA, EEF1A1, and AHNAK proteins. These were previously described as commonly identified BioID background proteins^48,49^, typically found to be highly abundant in BioID-only samples. For all proteins in both data sets, the iBAQ intensities were either significantly higher in the T2A condition or were not significantly different (Fig. 6a). This shows the background in our data sets are consistent with the BioID-only background observed in classical BioID experiments. For NCAP, we did not identify TOP1 in any of the T2A or MUTT2A samples. To extend observations beyond these few proteins, we integrated the BioID CRAPome^50^. The CRAPome was filtered to contain proteins that are identified in at least 25 out of 30 experiments with an average spectral count of 3 across all BioID experiments. This provided us with 382 proteins that are consistently identified in BioID experiments. Of note, of the aforementioned proteins TOP1 and PRKDC did not make it into this list, due to being identified in 16 and 23 out of 30 experiments, respectively. In addition, TOP1 had an average spectral count of 2.75 across all experiments, below our cutoff of 3. We then ranked our data sets based on enrichment and assessed the distribution of these 382 proteins within our data sets. For all data sets, most CRAPome proteins were distributed around a LOG2FC of 0, indicating that these proteins reside within the stable background (Fig. 6b). The data demonstrates that our T2A control allows efficient filtering of the same stable background as seen in other BioID experiments.

**Fig. 6.**
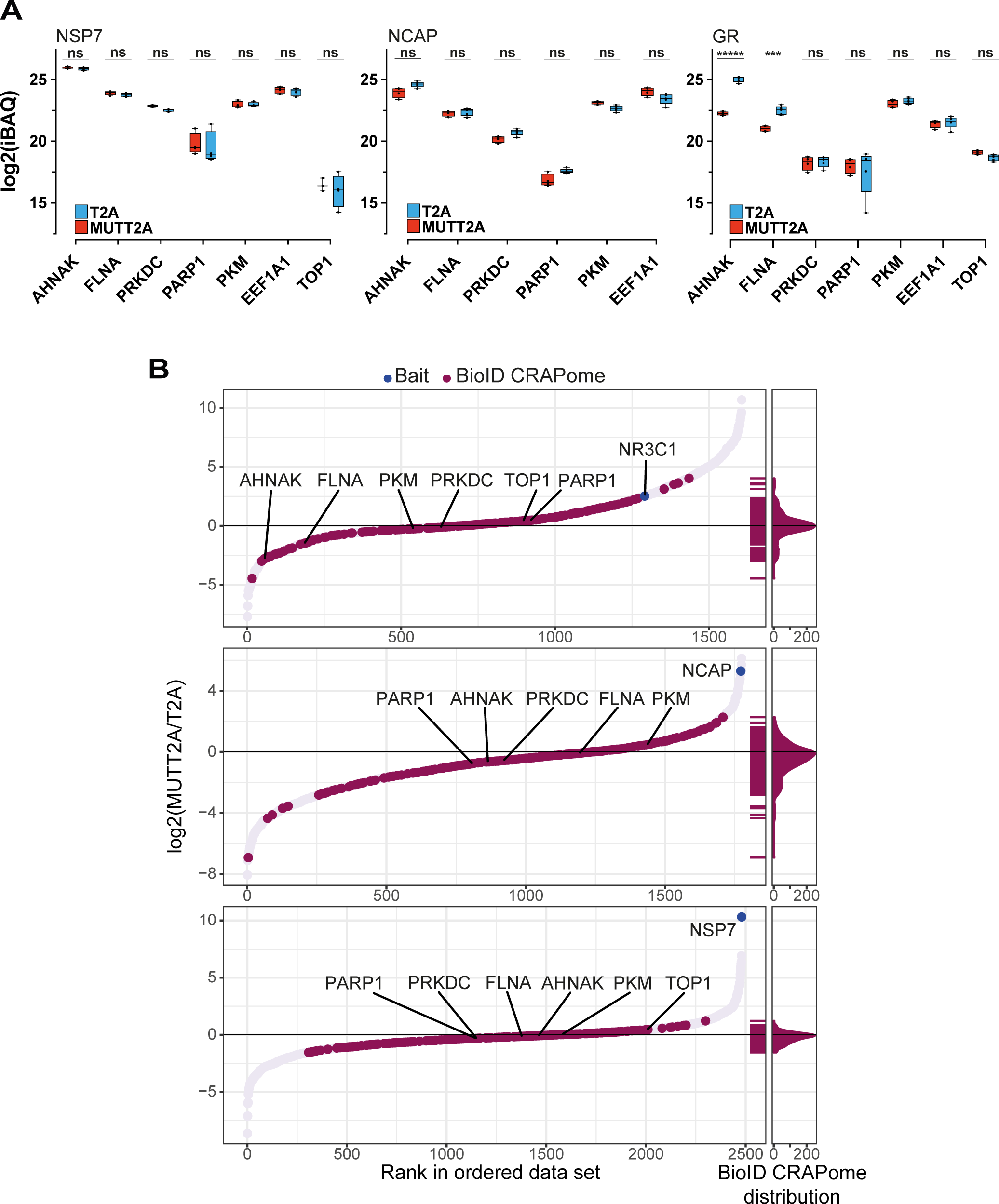
T2A split-link experiments demonstrate a similar background as classical BioID experiments. (a) iBAQ intensities of AHNAK, FLNA, PKM, PRKDC, TOP1, and PARP1 in each experiment. (b) Distribution of the BioID CRAPome within each experiment. Baits are shown in blue, proteins shared with the BioID CRAPome are shown in crimson. ***, adj. P-value < 0.05; *****, adj. P-value < 0.0005.

## DISCUSSION

Proximity labeling proteomics provides a powerful approach to identify protein-protein interactions in a wide variety of both *in vitro* and *in vivo* settings. However, only a limited amount of studies have sought to expand the toolbox beyond classically-used Flp-In 293T-Rex cells. Yet to expand proximity labeling to more relevant cellular settings, other integration methods are preferred as Flp-In requires pre-engineered cell lines which can be tedious to develop. Although alternative integration methods have been explored previously^42,51^, these studies, understandably, built BioID-compatible vectors for their own experimental questions based on classical ‘copy-paste’ cloning efforts. To accommodate these shortcomings, Samavarchi-Tehrani *et al.* ^42^ built a set of lentiviral transfer vectors that allow N- and C-terminal tagging of the protein-of-interest (POI) with BirA*. Although these vectors are a major advancement in the field, they lack the possibility of a customizable design and assembly. Haldeman et al. ^52^ built a Gateway^TM^-based modular cloning system that encompasses BioID2 and APEX labeling enzymes. However, inherent to Gateway^TM^ cloning are large scar sequences that cannot be omitted, the limited number of components that can be assembled, and the relatively expensive cost of recombinase reagents. These restrictions hamper throughput and versatility. Therefore, inspired by efforts such as MoClo^53^, here, we built a two-step Golden Gate-based toolbox for full custom assembly of BioID vectors, not only allowing to choose promoters, orientation of the proximity labeling enzyme to the POI, positive selection markers, etc., but also allowing to choose the type of vector backbone depending on the envisioned application. Although here we only present lentiviral and PB transposon vector backbones, the system can easily be expanded to accommodate vectors for other types of delivery or integration (e.g. AAV, <ΔC31 integrase, Sleeping Beauty, Flp-In) as well as for other organisms.

Today, proximity labeling proteomics acts as a complementary method to classical affinity purification for the identification of protein interaction partners, yet little effort has been made to optimize negative controls. This is surprising as a suitable negative control is a prerequisite to do an adequate quantitative differential analysis. Here, we describe the use of a T2A self-cleaving peptide as a suitable negative control. By introducing a single point mutation, we engineered a mutant T2A (MUTT2A) that no longer retains the capacity to cause ribosomal skipping. Recently, Sears *et al.* ^48^ reiterated good practices for BioID experiments, which includes a BioID-only control to filter for stochastic background interactions. Our T2A split-link extends upon the BioID-only control and allows to co-express the biotin ligase and the POI at equimolar amounts yet physically separated. As such, (quantitative) proteomic changes because of POI (or biotin ligase) overexpression remain present, while TurboID and POI levels can be evenly matched between the T2A and MUTT2A setup simply by optimizing doxycycline amounts for both cell lines. Our results demonstrate the validity of the T2A split-link as we identify known interactors for three different bait proteins. Notably, Chojnowski et al. ^54^ developed a conceptually similar method called 2C-BioID in which the biotin ligase and POI are tagged with either part of the chemically inducible FKBP-FRB oligomerization system. Upon supplementation of rapamycin or a biologically inactive rapalog, the biotin ligase is recruited to the POI, and differential analysis is performed by comparing supplemented and non-supplemented conditions. As such the biotin ligase itself does not interfere with proper POI localization. Moreover, the same cell line can be used for both conditions, whereas our T2A split-link needs at least two engineered cell lines. Nonetheless, both methods complement each other’s flaws. While 2C-BioID mainly addresses localization issues, T2A split-link does not require a chemical supplement which can perturb the behavior of the cell nor does it require the biotin ligase to be recruited. We demonstrate that differential analysis of MUTT2A compared to T2A generates a stable background as evidenced by the representation of proteins that are known to be highly abundant in BioID experiments. Interestingly, TurboID itself was always significantly enriched in the MUTT2A condition, which we also observed previously in our endogenous p53-(MUT)T2A-BioID^10^. Although seemingly counterintuitive, the difference in bait size between the MUTT2A and T2A setup very likely explains these observations. As the fusion protein in the MUTT2A setup represents an amount of biotinylatable amino acids that is larger compared to TurboID alone, as in the T2A setup, a higher degree of self-biotinylation and thus enrichment would be expected in the MUTT2A samples. This would be consistent with the significant yet overall modest enrichment of TurboID in our NSP7 data compared to the other POIs, as NSP7’s small size would leave room for only a limited amount of biotinylatable amino acids.

For NCAP and NSP7, we noticed the number of differential proteins was skewed in the T2A condition compared to the MUTT2A. We reasoned this to be due to a differential localization of the biotin ligase. Indeed, we observed a higher amount of nuclear proteins in the T2A condition, likely representing that free TurboID can diffuse into the nucleus, something that was already shown for other BirA-derived biotin ligases. Here, we show this to also be the case for TurboID. The impact of this differential localization on the number of known interactors, however, seems to be relatively limited.

Taken together, we provide the interactomics community with a versatile platform for the generation of proximity ligation tools in a variety of vector backbones. In addition, we expand the proximity ligation toolbox with a more suitable negative control compared to previously published options. We demonstrate our T2A split-link approach compares nicely with published data for three different bait proteins. Finally, we show that our differential analysis of our T2A split-link concept allows efficient filtering of commonly observed BioID contaminants.

## STAR METHODS

### Resource availability

#### Lead contact

Requests for resources and additional information should be directed to and will be fulfilled by the lead contact, Dr. Sven Eyckerman.

#### Materials availability

Plasmids generated in this study are available via the BCCM/GeneCorner Plasmid Collection (genecorner.ugent.be) with accession numbers 13830 to 13835.

#### Data and code availability

Proteomics data have been deposited to the ProteomeXchange Consortium via the PRIDE partner repository and are publicly available as of the data of publication. Accession numbers and reviewer log in data are listed in the key resources table. Uncropped western blots are found in Fig. S6. Microscopy data reported in this paper will be shared by the lead contact upon request. This paper does not report original code. Any additional information required to reanalyze the data reported in this paper is available from the lead contact upon request.

### Experimental model and subject details

#### Cell lines

Human HEK293T and A549 cell lines were cultured in DulBecco’s Modified Eagle Medium (DMEM) supplemented with 10% FBS. Human SK-N-BE(2)-C and SHEP neuroblastoma cell lines were cultured in Roswell Park Memorial Institute (RPMI) 1640 medium supplemented with 10% FBS. Parental cell lines were maintained in antibiotic-free conditions, experiments and transductions were performed with 30 U/mL Penicillin-Streptomycin. Cells were kept under 60-70% confluency and passaged twice a week. Cells lines were confirmed mycoplasma-free by a mycoplasma PCR detection kit. Transgenic cell lines were maintained in the same medium as the parental lines but continuously supplemented with 30 U/mL Penicillin-Streptomycin. Transgenic lines were regularly pulsed with the appropriate antibiotic for the corresponding transgene. Cells were maintained at 5% CO_2_ on 37°C.

### Method details

#### Molecular cloning

All backbones contained a dual selection cassette between the BsaI/BsmBI overhangs expressing a ccdB toxin and a chloramphenicol (*cat*) gene, conferring negative and positive selecting respectively. These plasmids were propagated in 2T1R, XL-10 Gold or DB3.1 competent cells which either contain a *gyrA* R462C conversion or the F’ plasmid expressing the ccdA antitoxin, making them resistant to ccdB negative selection.

To generate level-0 Golden Gate modules, all parts were either cloned by PCR or DNA synthesis with BsaI sites that generate the corresponding overhangs required for the module. Parts were mixed with their corresponding module vector (pGGAB, pGGBC, pGGCD, pGGDE, pGGEF, pGGFG) at a 3:1 ratio (w:w) with a minimum of 50 ng of module vector. These mixes were digested with BsaI-HFv2 for 1 h at 37°C in 1X CutSmart buffer. After digestion, the reaction was stopped by heating to 80°C for 20 min. Reactions were cooled to room temperature, and 1X T4 ligase buffer and 1 uL T4 DNA ligase was spiked in the reaction mixture. Ligation was performed for at least 1 h up to overnight incubation at room temperature. Ten microliters of the reaction mixture were chemically transformed in DH10B and selected on LB agar plates containing 50 ug mL^−1^ carbenicillin. Colonies were screened by colony PCR with GoTaq G2 master mix and a diagnostic restriction digest with EcoRI and HindIII. Positive clones were sequence-verified by Sanger sequencing.

Level-1 backbone plasmids were generated as indicated in the main text. Level-1 plasmids were assembled by combining 100 ng of each part with 100 ng of backbone, 1 mM ATP, 200 U T4 DNA ligase, 10 U BsaI-HFv2, and 1X CutSmart buffer in a total volume of 15 uL. Assemblies were cycled for 20 cycles for 2 min at 37°C and 2 min at 16°C, followed by 5 min at 50°C and finally 5 min at 80°C to stop the reaction. Ten microliters of the assembly were chemically transformed in DH10B competent cells and selected on LB agar plates containing 50 ug mL^−1^ kanamycin. Colonies were screened with a diagnostic restriction digest depending on the assembled parts.

Lentiviral and piggyBac backbone vectors were generated as indicated in the main text. For plasmids generated by GGW, 75 ng of at least two level-1 plasmids and one backbone plasmid with compatible attL/R gateway sites was combined with 1X LR recombinase, and TE buffer (pH 8.0) in a total volume of 10 uL. Mixtures were incubated overnight at 25°C. One microliter proteinase K was added for 10 min at 37°C to stop the reaction. For BsmBI-based Golden Gate assembly, 100 ng of at least two level-1 plasmids and one backbone plasmid was combined with 1X T4 DNA ligase buffer, 200 U T4 DNA ligase, and 10 U BsmBI-v2 in a total volume of 15 uL. Assemblies were incubated for 30 cycles with 2 min at 42°C and 2 min at 16°C, one cycle of 5 min at 60°C, and finally one cycle of 5 min at 80°C to stop the reaction. In both cases, 10 uL of the reaction were chemically transformed in DH10B or Stbl3 competent cells, and selected on LB agar plates containing 50 ug mL^−1^ carbenicillin. Clones were screened with a diagnostic restriction digest depending on the assembled parts.

#### Lentivirus production and transduction

For lentiviral productions, 6.5 × 10^6^ HEK293T cells were seeded for calcium phosphate transfection the next day. Medium was refreshed 30 min to 4 h prior to transfection. The DNA mixture comprised 24 ug of transfer plasmid, 18 ug pCMV-dR8.74, and 7.2 ug pMD2.G, 75 uL CaCl_2_ (2.5 M) in a total volume of 750 uL in sterile water. The DNA mixture was added dropwise to 750 uL HEPES-buffered saline (Sigma-Aldrich) while vortexing. The transfection mixture was incubated 5 min at room temperature, and added to the cells. Next day, a clear calcium phosphate precipitate was observed and the medium was refreshed to avoid toxicity of the transfection reagents. The following two days, each day medium containing lentiviral particles was harvested and kept at 4°C. After the last harvest, both harvests were combined and the medium was spinned at 500 x g for 5 min at 4°C to pellet cellular debris. Supernatant was filtered through a 0.45 um filter, and lentiviruses were pelleted by ultracentrifugation for 2.5 h at 85.000 x g at 4°C. Pellets were redissolved in 100 uL DMEM and aliquoted per 20 uL.

To transduce SHEP and SK-N-BE(2)-C neuroblastoma cells, 1 × 10^6^ cells were seeded. The day after seeding, 5 uL concentrated lentivirus was added to the medium. Next day, the medium was refreshed. Two days post-transduction, 4 and 1 ug mL^−1^ puromycin was added to the SK-N-BE(2)-C and SHEP cells, respectively. One million A549 cells were transduced with a multicomponent tetracycline-inducible transactivation system at a MOI of 3, expressing a transcriptional repressor tTS and a transactivator rtTA at equimolar amounts. tTS actively silences the TRE promoter in the absence of doxycycline but is displaced upon the addition of doxycycline. The rtTA transactivator acts oppositely when doxycycline is supplemented, actively transcribing genes inserted downstream of the TRE promoter region. Two days post-transduction, 400 ug mL^−1^ hygromycin was added. Optimal puromycin and hygromycin concentrations were determined prior by serial dilution of the antibiotic for the neuroblastoma cell lines and A549, respectively. Transduced cells were selected for a minimum of 2 days or 2 weeks for puromycin and hygromycin, respectively, and until the parental line (under the same selection regime) was no longer alive.

#### PiggyBac transposition

Twenty thousand A549 tetracycline-inducible transactivator cells were seeded in 500 uL in a 24-well plate for transfection the next day. A total amount of 500 ng plasmid DNA, comprising 400 ng TurboID plasmid and 100 ng pCMV-hyPBase (Wellcome Trust Sanger Institute^55^), was diluted to 100 uL with Opti-MEM, after which 0.5 uL PLUS reagent was added. The mixture was incubated for 10 min at room temperature. Next, 1.5 uL Lipofectamine LTX was added, and incubated for 25 min at room temperature after gently mixing. Medium was refreshed with 500 uL antibiotic-free medium, and the DNA-liposome mixture was added to the cells. Next day, the medium was refreshed to complete growth medium with antibiotics. One week post-transfection, 5 ug mL^−1^ blasticidin was supplemented to the growth medium for a minimum of 2 weeks, and until the parental line (under the same selection regime) was no longer alive. The optimal blasticidin concentration was determined prior by kill curve analysis with a serial dilution of the antibiotic.

#### SDS-PAGE and western blotting

Thirty micrograms of protein material was measured using Bradford reagent (Bio-Rad Protein Assay Dye Reagent concentrate #5000006). Each sample was supplemented with 7.5 uL XT Sample Buffer (Bio-Rad #1610791) and 1.5 uL XT Reducing Agent (Bio-Rad #1610792) in a total final volume of 30 uL. Samples were heated to 95°C for 10 min, cooled down prior to being loaded and ran on a 4-12% ExpressPlus PAGE 4-12% pre-cast gel (Genscript M421215) according to the manufacturer’s instructions. Proteins were transferred to PVDF membrane (Merck #IPFL00010) for 3 h at 60 V in methanol blotting buffer (48 mM Tris-HCl, 39 mM glycine, 0.0375% SDS (w:v), and 20% methanol (v:v)). Membranes were blocked for 30 min at room temperature with Odyssey Blocking buffer (LI-COR 927-50000) Primary antibodies were incubated overnight at 4°C with gentle end-to-end rotation. The following primary antibodies and dilutions in TBS were used: mouse monoclonal anti-V5 antibody (Invitrogen R960-25) at 1/5000 and rabbit polyclonal anti-ACTB (Sigma-Aldrich #A2066) at 1/2000. Membranes were washed three times with TBST for 10 min at room temperature. The following secondary antibodies were used at a dilution of 1/5000: goat polyclonal anti-mouse IgG IRDye 800CW (LI-COR), goat polyclonal anti-rabbit IgG IRDye 680RD (LI-COR). Membranes were incubated with secondary antibodies for 1 h at room temperature, washed three times in TBST, and visualized on a LI-COR Odyssey IR scanner. For streptavidin staining, membranes were washed three times with TBST after visualization and incubated for 1 h at room temperature with IRDye 680RD Streptavidin (LI-COR) at a 1/50000 dilution. After incubation, membranes were washed once more and biotinylated proteins were visualized be re-scanning the membrane.

#### RNA extraction, cDNA synthesis and RT-qPCR

Per condition, 1 × 10^6^ cells per well were plated in a 6-well plate. One day after plating, cells were incubated with the optimized doxycycline concentrations depending on the cell line as indicated in the main text. Twenty-four hours post-induction cells were lysed on the plate directly by addition of 350 uL RA1 (Macherey-Nagel, 740961) supplemented with 3.5 uL β-mercaptoethanol. RNA was extracted from the mixture using the Nucleospin RNA mini kit following the manufacturer’s instructions (Macherey-Nagel, 740955). RNA was eluted from the column in 80 uL RNase-free water, and RNA concentration and quality was assessed spectrophotometrically. Two hundred fifty nanograms of RNA was used for cDNA synthesis using the PrimeScript RT kit (Takara Bio, RR037A). Primers listed in Table S4 were used to amplify housekeeping genes (SDHA, YWHAZ, and UBC) and targets-of-interest (TSC22D3 and DUSP1). qPCRs were performed using the SensiFAST SYBR No-ROX kit (Meridian Bioscience, BIO-98005) consisting of 0.5 uL 10 uM of each primer, 5 uL 2X SensiFAST SYBR No-ROX mix, and 12.5 ng cDNA. Samples were measured in technical duplicates. Fluorescent signal was detected using a LightCycler 480 System (Roche). The following cycling conditions were used: 1 cycle at 95°C for 5 min, 40 cycles at 95°C for 10 s, 60°C for 10 s and 72°C for 10 s, followed by melting curve analysis. Quantitation cycles (Cq) of target genes were normalized to all three housekeeping genes using geometric averaging^56^. The geNorm algorithm was used to calculate stability of housekeeping genes across samples. All RT-qPCR analyses were performed in qbase+ (CellCarta).

#### Proximity labeling

For each proximity labeling experiment, three 150 cm^2^ plates were seeded with 2.7 × 10^6^ cells per replicate. Next day, optimized doxycycline concentrations as previously described were added to the corresponding setup. After 24 h, 50 uM biotin was added for 1 h. Cells were washed once on the plate with 10 mL ice-cold PBS, after wish cells were collected per replicate by scraping in 750 uL ice-cold PBS. Cells were pelleted by spinning for 5 min at 4°C at 500 x g. Pellets were washed once with 6 mL ice-cold PBS. Pellets were frozen at −20°C until further processing. Subsequently, cell pellets were resuspended in 5 mL ice-cold RIPA (50 mM Tris-HCl pH 7.5, 150 mM NaCl, 1% NP-40, 2 mM EDTA, and 0.1% SDS in ddH_2_O), supplemented with 1X cOmplete Protease Inhibitor cocktail and 0.5% sodium deoxycholate. Next, 50 U mL^−1^ benzonase was added and samples were incubated for 1 h at 4°C with end-to-end rotation. Lysates were then sonicated on ice with a probe sonicated at 30% amplitude for 5 rounds of 6 s burst with 2 s in between rounds. Lysates were pelleted at 16100 x g for 15 min at 4°C and the supernatant was transferred to a new tube. Protein concentrations were determined by Bradford assay, and a maximal shared protein amount across all samples in the same experiment was used as an input for the affinity precipitation. Per replicate 30 uL Streptavidin Sepharose High Performance beads were washed three times with 600 uL unsupplemented ice-cold RIPA and eventually resuspended in 600 uL ice-cold supplemented RIPA buffer. Samples volumes were adjust to a minimum volume of 4.5 mL, and equilibrated beads were added to each sample. Affinity purification was performed by incubation at 4°C for 3 h with end-to-end rotation. Next, beads were pelleted by centrifugation for 1 min at 500 x g. Beads were washed with 1 mL unsupplemented RIPA buffer. This process was repeated for a total of three washes, after which beads were washed twice with freshly prepared ABC buffer (50 mM NH_4_HCO_3_ pH 8.0 in ddH_2_O). Next, beads were washed three times with trypsin digest buffer (20 mM Tris-HCl pH 8.0 and 2 mM CaCl_2_ in ddH_2_O). Ultimately, beads were resuspended in 20 uL Tris-HCl pH 8.0, and 1 ug trypsin was added. Samples were incubated overnight at 37°C. Next morning, beads were pelleted and supernatant was transferred to a new tube. Another 0.5 ug trypsin was added and samples were incubated for an additional 3 h at 37°C. Peptide mixtures were acidified with 20% FA to a final concentration of 2% FA. Mixtures were centrifuged at 20000 x g for 10 min at room temperature, and the supernatant was transferred to a MS vial. Samples were frozen at −20°C until LC-MS/MS analysis.

#### Confocal imaging

Ten thousand cells per chamber were seeded in an 8-chamber imaging slide. Next day, optimized doxycycline concentrations were added for 24 h. Two hours prior to fixation 1 uM dexamethasone was added to T2A-GR and MUTT2A-GR cell lines, and 1 h prior to fixation 50 uM biotin was added. Half in each chamber was removed and replaced with the same volume of 4% PFA in PBS for a final concentration of 2% PFA. PFA was pre-warmed to 37°C prior to use. Cells were incubated at room temperature for 15 min with gentle horizontal shaking. Subsequently, cells were gently washed three times with PBS, followed by three washing steps with 0.2% Triton X-100 in PBS for 5 min each to permeabilize the cells. After three more washes with PBS, cells were blocked for 1 h at room temperature with blocking buffer (0.5% BSA, 0.02% Triton X-100, 1:100-diluted donkey serum in PBS). For V5 staining, cells were incubated overnight with 1:500-diluted mouse anti-V5 primary antibody in blocking buffer. Next day, cells were washed three times with PBS before being incubated for 2 h with donkey anti-mouse AlexaFluor 568 secondary antibody and 1 h with streptavidin DyLight 488. After three PBS washes, cells were stained with 1:1000-diluted DAP for 15 min at room temperature to stain nuclei. Cells were washed three more times with PBS and were kept in PBS until confocal imaging. All imaging was performed on a LSM880 Airyscan with a Plan-Apochromat 63x/1.4 Oil DIC M27. The Airyscan detector was operated in the super resolution mode of the FastAiryScan. In ZEN Black 2.3 SP1, a pixel reassignment and 2D wiener deconvolution were carried out post-acquisition. Image processing was performed in ZEN Blue 3.5 or Fiji.

#### LC-MS/MS and data analysis

For each sample, 2.5 uL peptide mixture was injected for LC-MS/MS analysis on a Ultimate 3000 RSLCnano system (Thermo Fisher Scientific) in line connected to a Q-Exactive-HF Biopharma mass spectrometer (Thermo Fisher Scientific). Trapping was performed 20 uL min^−1^ for 2 min in loading solvent A (98% ACN, 0.1% TFA) on a 5 mM trapping column (Thermi Fisher Scientific). Peptide separation was performed on a 250 mm Aurora Ultimate (IonOpticks) at a constant temperature of 45°C. Peptides were eluted by a non-linear gradient starting at 1% solvent B (80 ACN, 0.1% FA) reaching 33% solvent B in 60 min, 55% in 75 min, 70% in 90 min, followed by a wash at 70% solvent B for 10 min and re-equilibration with solvent A. The mass spectromter was operated in a data-dependent acquisition mode, automatically switching between MS1 and MS2 acquisition for the 12 most abundant ion peaks per MS1 spectrum. Full-scan MS spectra (375 – 1500 m/z) were acquired at a resolution of 60000 in the Orbitrap analyzer after accumulation to a target value of 3000000. The 12 most intense ions above a threshold of 15000 were isolated for fragmentation at a normalized collision energy of 30%. The C-trap was filled at a target value of 100000 for maximum 80 ms and the MS2 spectra (200 – 2000 m/z) were acquired at a resolution of 15000 in the Orbitrap analyzer with a fixed mass of 145 m/z. Only peptides with charge states ranging from +2 to +6 were included for fragmentation and the dynamic exclusion was set to 12 s.

RAW files were searched using the Andromeda search engine with default search settings (1% FDR at peptide and protein level) as implemented in MaxQuant (v2.3.4.0). Spectra were searched against the human SwissProt proteome database (version of January 2023). Sequences of V5-TurboID, SARS-CoV-2 NCAP, and SARS-CoV-2 NSP7 were added to the search database. Enzyme specificity was set as C-terminal to arginine and lysine (trypsin), also when followed by a proline, with a maximum of two missed cleavages. Methionine oxidation and N-terminal acetylation were set as variable modifications. No fixed modifications were set. Matching between runs was disabled. Only proteins with at least one unique or razor peptide were retained for identification. Proteins were quantified using the MaxLFQ algorithm with iBAQ turned on.

Proteins that are known contaminants, identified as reverse hits, or only identified by site, were removed from the analysis. iBAQ values were log2-transformed and each replicate was median-normalized. Based on PCA sample clustering, outlying replicates were removed from the analysis. Proteins only identified in N-1 replicates in either the T2A or MUTT2A condition were retained for downstream statistical analysis, missing values were imputed by quantile regression with the imputeLCMD R package. In limma, a linear model was fitted onto the data and differential analysis was performed by empirical Bayes moderated t-tests with a Benjamini Hochberg correction for multiple testing. Proteins with an adjusted p-value of less than or equal to 5% were considered statistically significant.

#### Gene set enrichment analysis

All GSEA analyses were ran pre-ranked including all quantifiable proteins in the data set. These were ranked on the basis of the log2 fold change of their iBAQ values in the MUTT2A versus T2A condition. For statistical evaluation, the analyses were ran with 10000 permutations and an FDR correction. GSEA was performed with the curated gene sets (C2, v2023.1) and ontology gene sets (C5, v2023.1) human collections within the Molecular Signatures Database. GMT files for Dendoncker *et al*.^24^, Lempiaeinen *et al*.^23^, Laurent *et al*.^35^, Liu *et al*.^31^, May *et al*.^33^, Meyers *et al*.^32^, and Samavarchi-Tehrani *et al.*^34^ were custom-made based on their data entries in BioGRID (v.4.4) for NR3C1, SARS-CoV-2 NCAP, and SARS-CoV-2 NSP7. GSEA analyses and visualization were performed using the GSEA function within the clusterProfiler^57^ R package.

#### Human Protein Atlas subcellular compartment

To assess protein subcellular localization, significantly enriched proteins in either the T2A or MUTT2A setups were queried against the immunofluorescence data of the Human Protein Atlas using the HPAanalyze^58^ R package. Proteins assigned to the terms ‘Nucleoplasm’ or ‘Cytosol’ were used to demonstrate overall enrichment of nuclear or cytosolic proteins in either setup.

### Key resource table

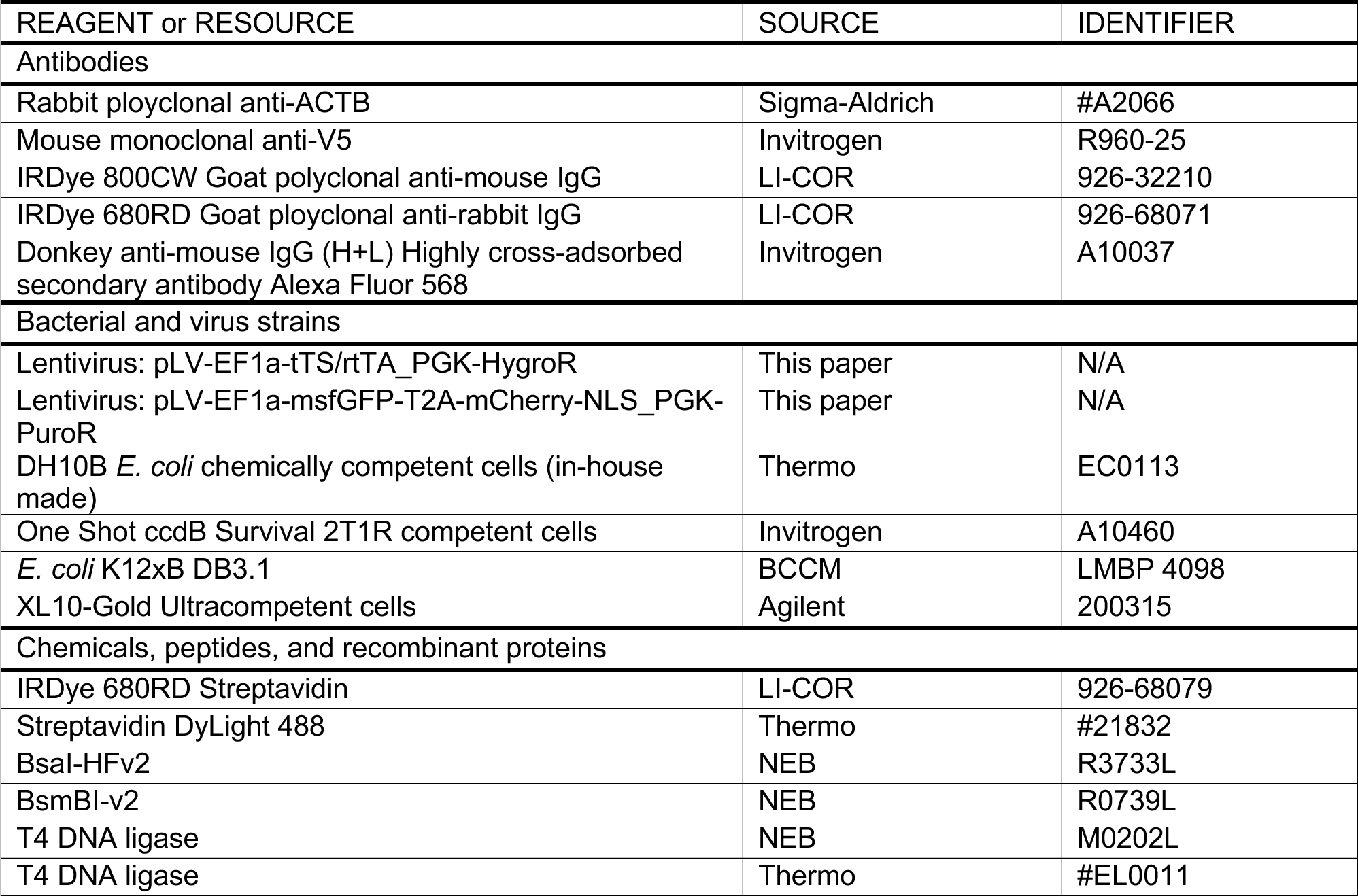
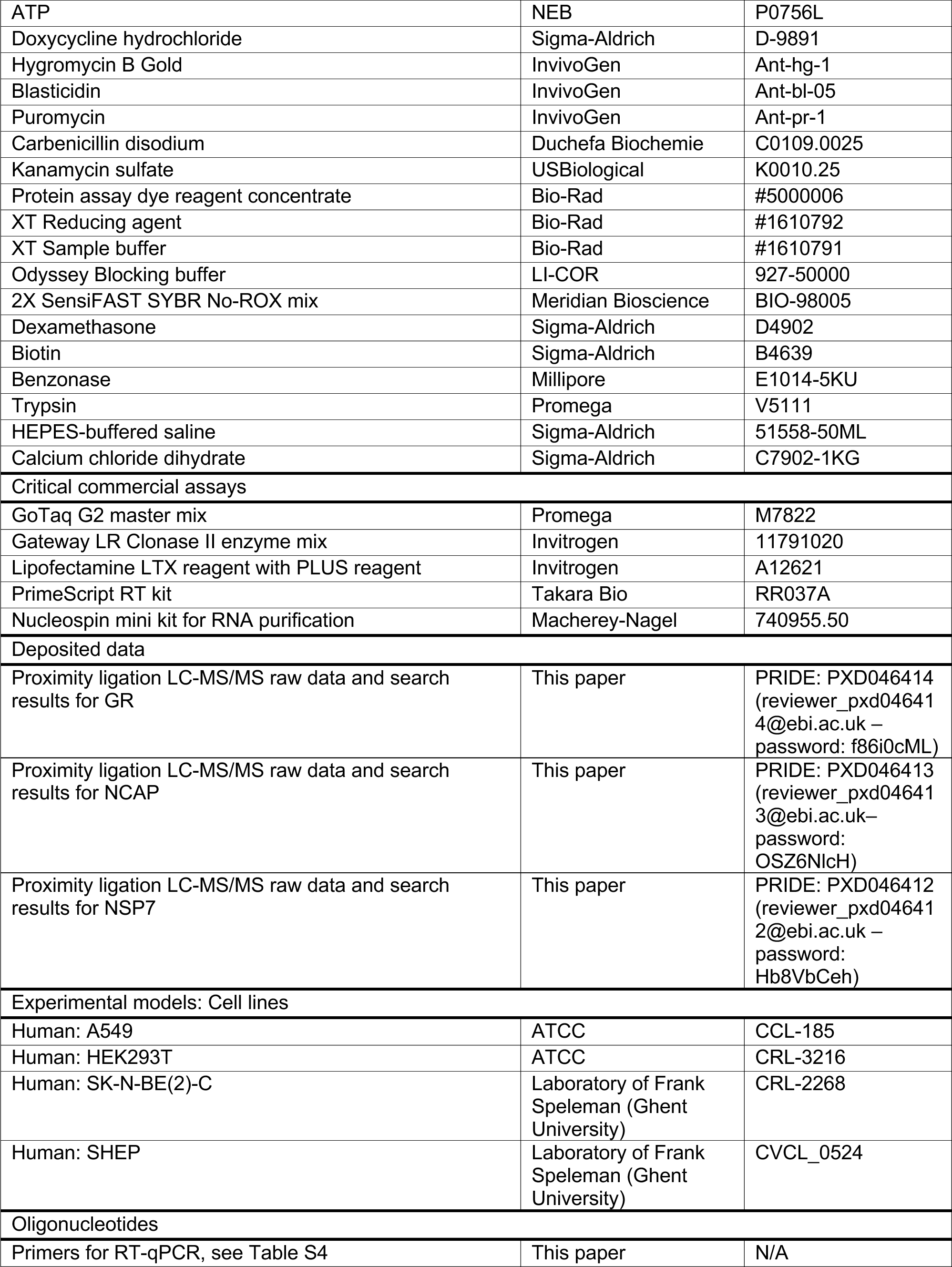
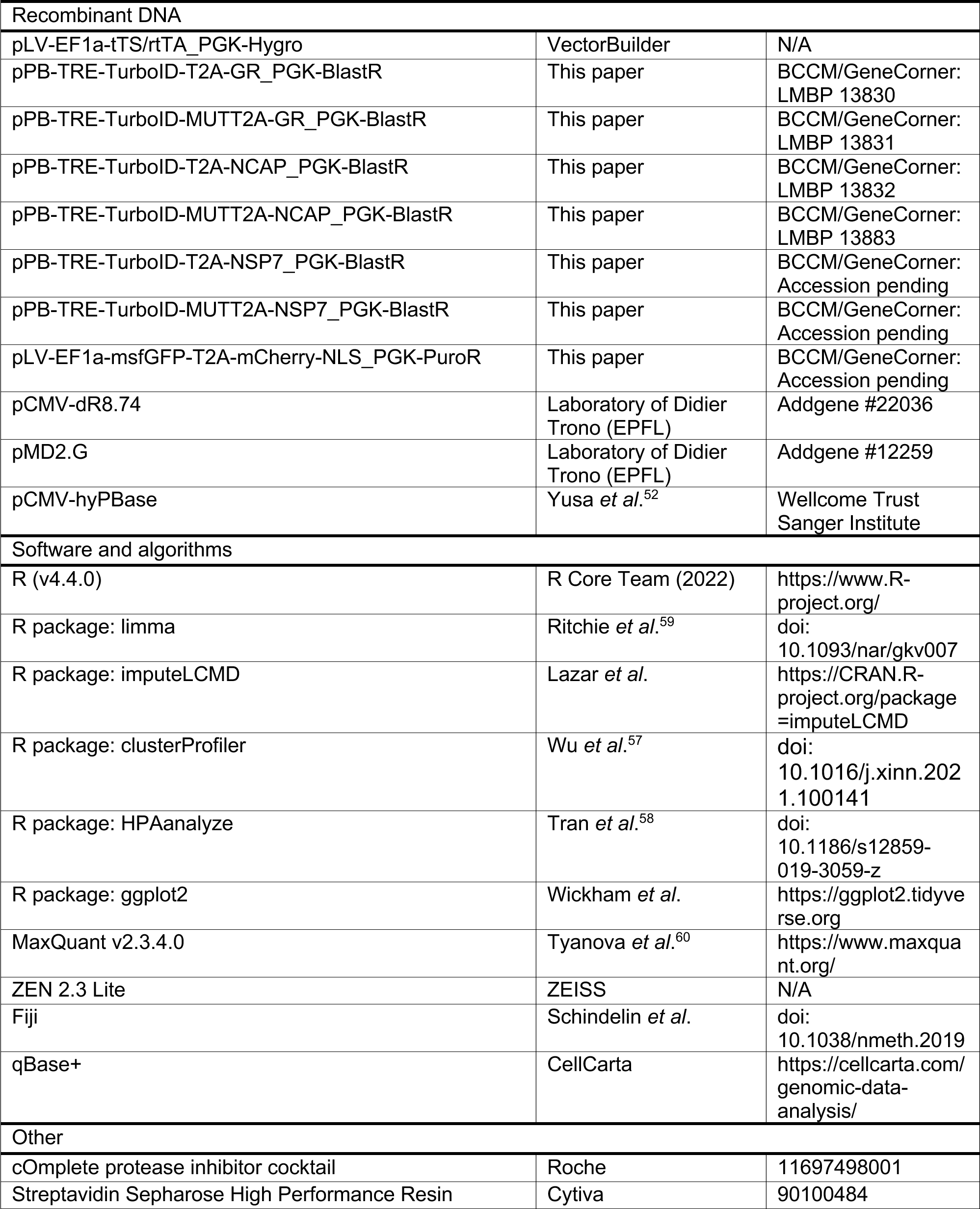

## Acknowledgements

The authors would like to acknowledge the VIB Proteomics Core and VIB Bioimaging Core Ghent for their assistance of the work presented in the manuscript. The authors would like to acknowledge funding support by a UGent BOF post-doctoral mandate to LD (BOF22/PDO/024), FWO PhD SB fellowships to GDM and LVM, and FWO (G042918N) and UGent BOF (BOF.GOA.2022.0003.03) projects to SE. The authors declare no competing interests.

## Author contributions

LD, GDM, AV, DDS, TVS, and MDM performed experimental work. LD, AV, DDS, and TVS performed molecular cloning. AV, DDS, and TVS performed BioID and LC-MS/MS sample preparation. LD and DF performed LC-MS/MS data analysis. LD and GDM performed lentivirus production and transduction. GDM and MDM performed and analyzed imaging data. LD performed and analyzed RT-qPCR. LVM, KDB, and TJ provided resources and support. LD, GDM, MDM, LVM, and SE did experimental design. LD drafted the first version of the manuscript and figures. All authors read the final manuscript and provided feedback. SE supervised and conceived the project.

---

**Fig. S1.**
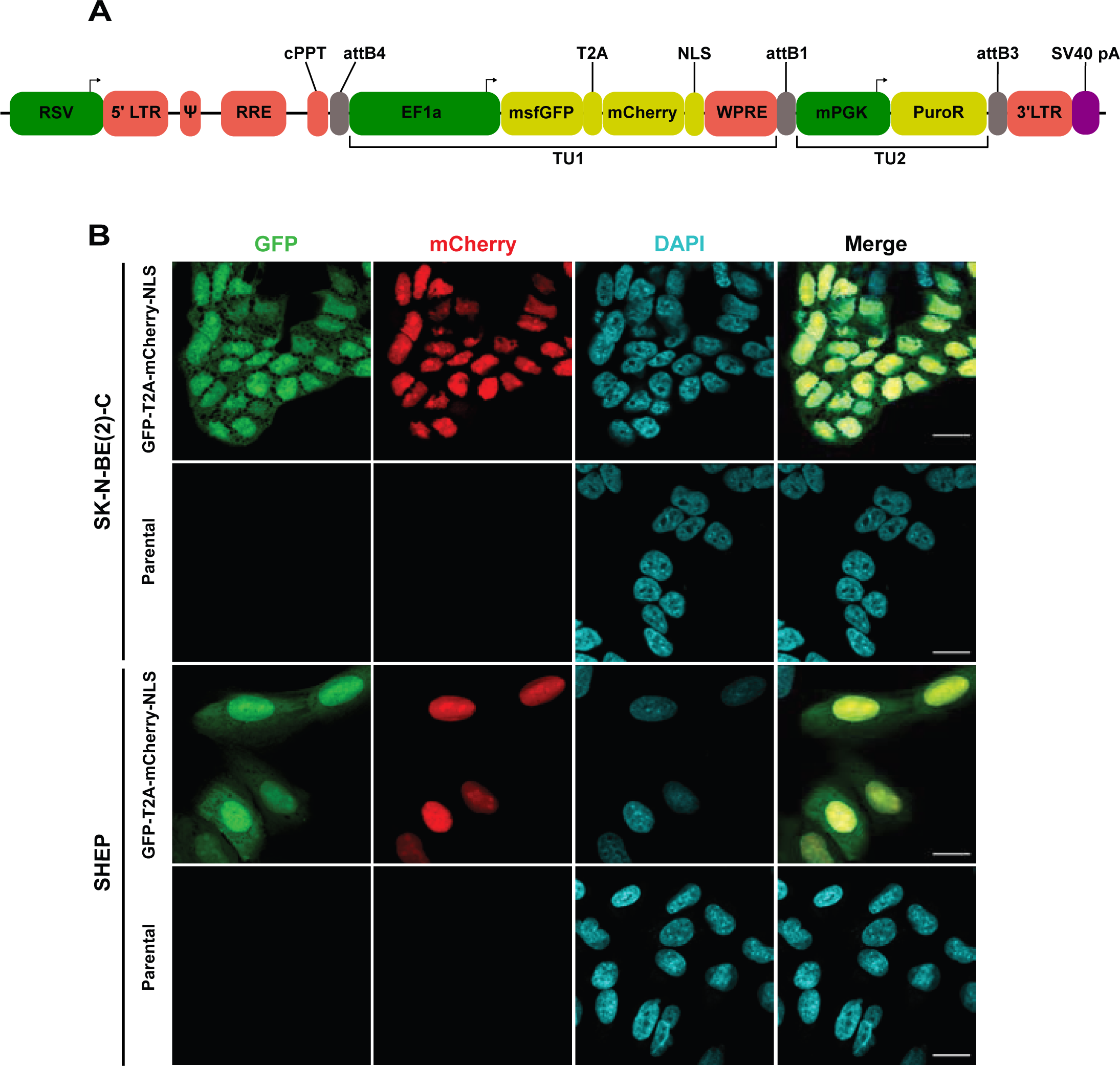
Golden GateWAY for assembly of mammalian expression constructs. (a) schematic overview of a proof-of-concept lentiviral construct expressing msfGFP and nuclear mCherry proteins. (b) Transduced SKNBE2C and SHEP cells expressing the proof-of-concept construct after puromycin selection. Parental cell lines are shown as negative controls for the immunofluorescence. RSV, respiratory syncytial virus promoter; LTR, long terminal repeat; RRE, Rev response element; cPPT, central polypurine tract; EF1a, elongation factor 1 alfa promoter; msfGFP, monomeric superfolder green fluorescent protein; NLS, nuclear localization signal; WPRE, Woodchuck Hepatitis Virus posttranscriptional regulatory element; mPGK, murine phosphoglycerate kinase 1 promoter; PuroR, puromycin resistance gene; pA, polyadenylation signal; TU, transcriptional unit.

**Fig. S2.**
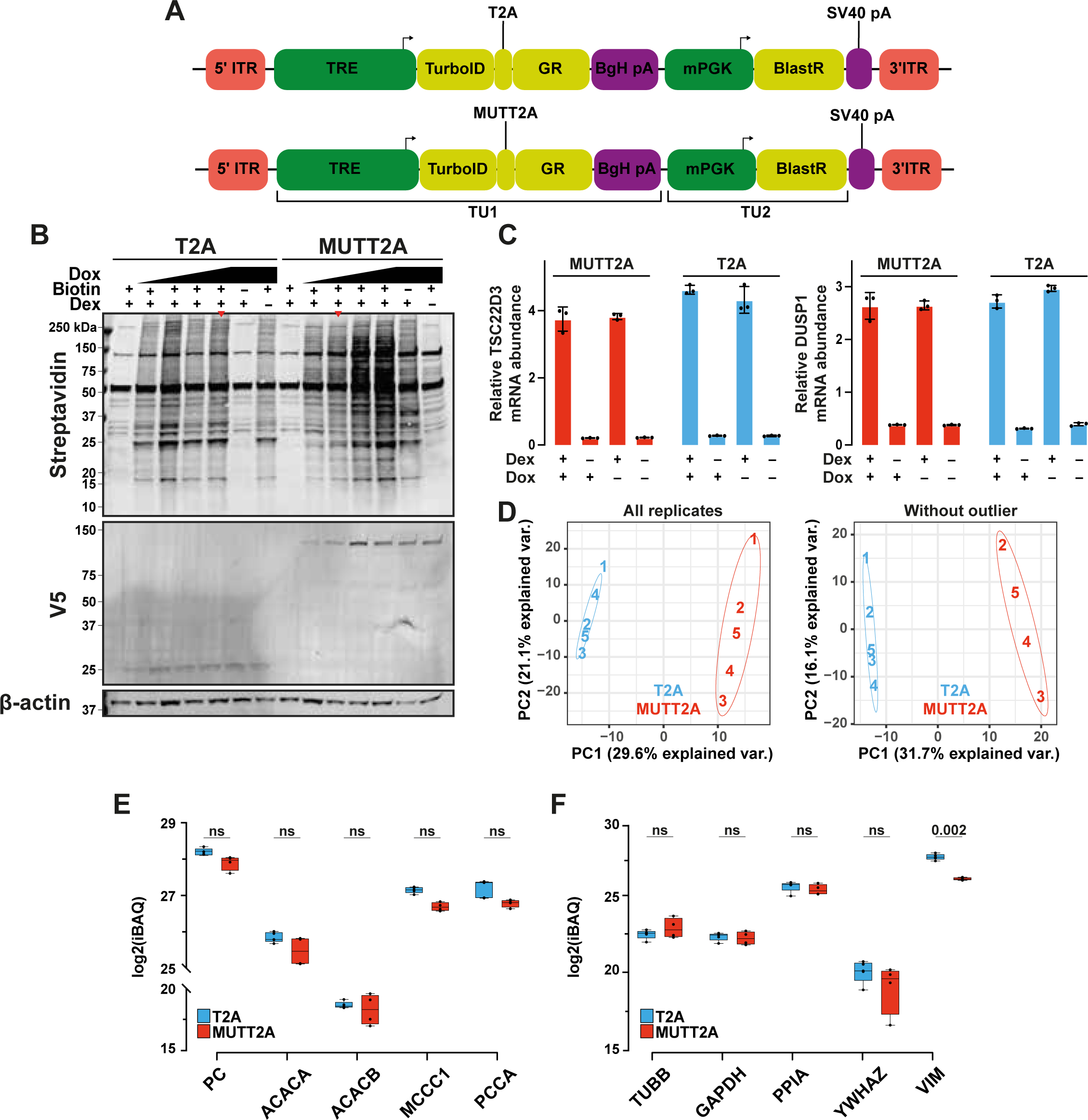
Engineering and validation of TurboID-T2A/MUTT2A-GR A549 cell lines. (a) Schematic overview of TurboID-T2A/MUTT2A-GR piggyBac transposon constructs. (b) Expression (V5) and biotinylation (streptavidin) staining of induced TurboID-T2A/MUTT2A-GR A549 cell lines. Red arrows indicate the doxycycline conditions used for downstream experiments. (c) mRNA expression of GR-target genes *TSC22D3* and *DUSP1*. (d) Principal component analysis before and after removing outlying replicates. (e) iBAQ intensities of endogenously biotinylated proteins. (f) iBAQ intensities of housekeeping proteins. ITR, inverted terminal repeat; TRE, tetracycline responsive element; GR, glucocorticoid receptor; BgH pA, Bovine growth hormone polyadenylation signal; mPGK, murine phosphoglycerate kinase 1 promoter; BlastR, blasticidine resistance gene; TU, transcriptional unit; Dox, doxycycline; Dex, dexamethasone; PC, principal component; iBAQ, intensity-based absolute quantification.

**Fig. S3.**
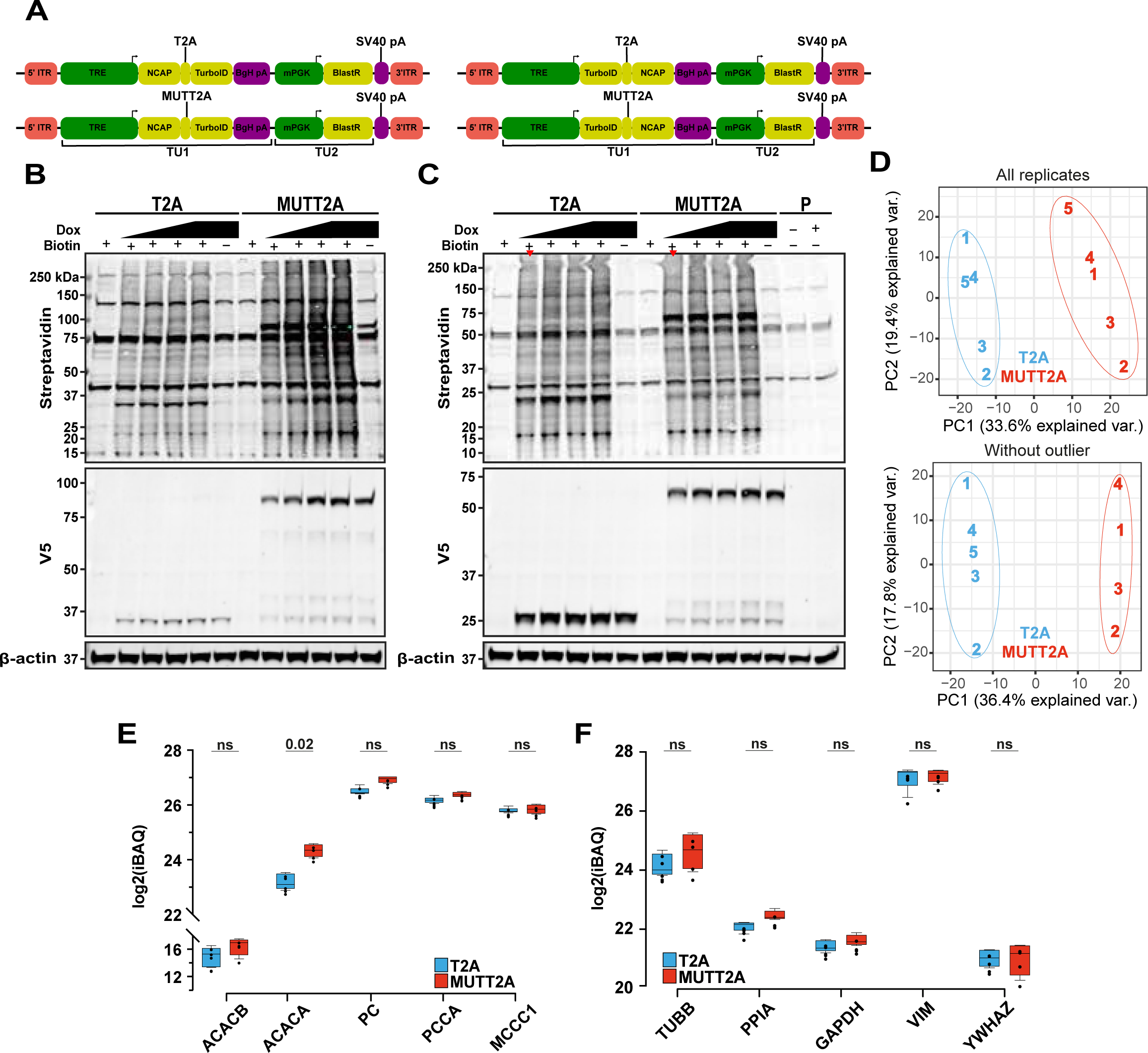
Engineering and validation of TurboID-T2A/MUTT2A-NCAP A549 cell lines. (a) Schematic overview of N- and C-terminally tagged NCAP with T2A/MUTT2A-TurboID piggyBac transposon constructs. (b,c) Expression (V5) and biotinylation (streptavidin) staining of A549 cells engineered with constructs shown in (a). (d) Principal component analysis before and after removing outlying replicates. (e) iBAQ intensities of endogenously biotinylated proteins. (f) iBAQ intensities of housekeeping proteins. ITR, inverted terminal repeat; TRE, tetracycline responsive element; GR, glucocorticoid receptor; BgH pA, Bovine growth hormone polyadenylation signal; mPGK, murine phosphoglycerate kinase 1 promoter; BlastR, blasticidine resistance gene; TU, transcriptional unit; Dox, doxycycline; Dex, dexamethasone; PC, principal component; iBAQ, intensity-based absolute quantification.

**Fig. S4.**
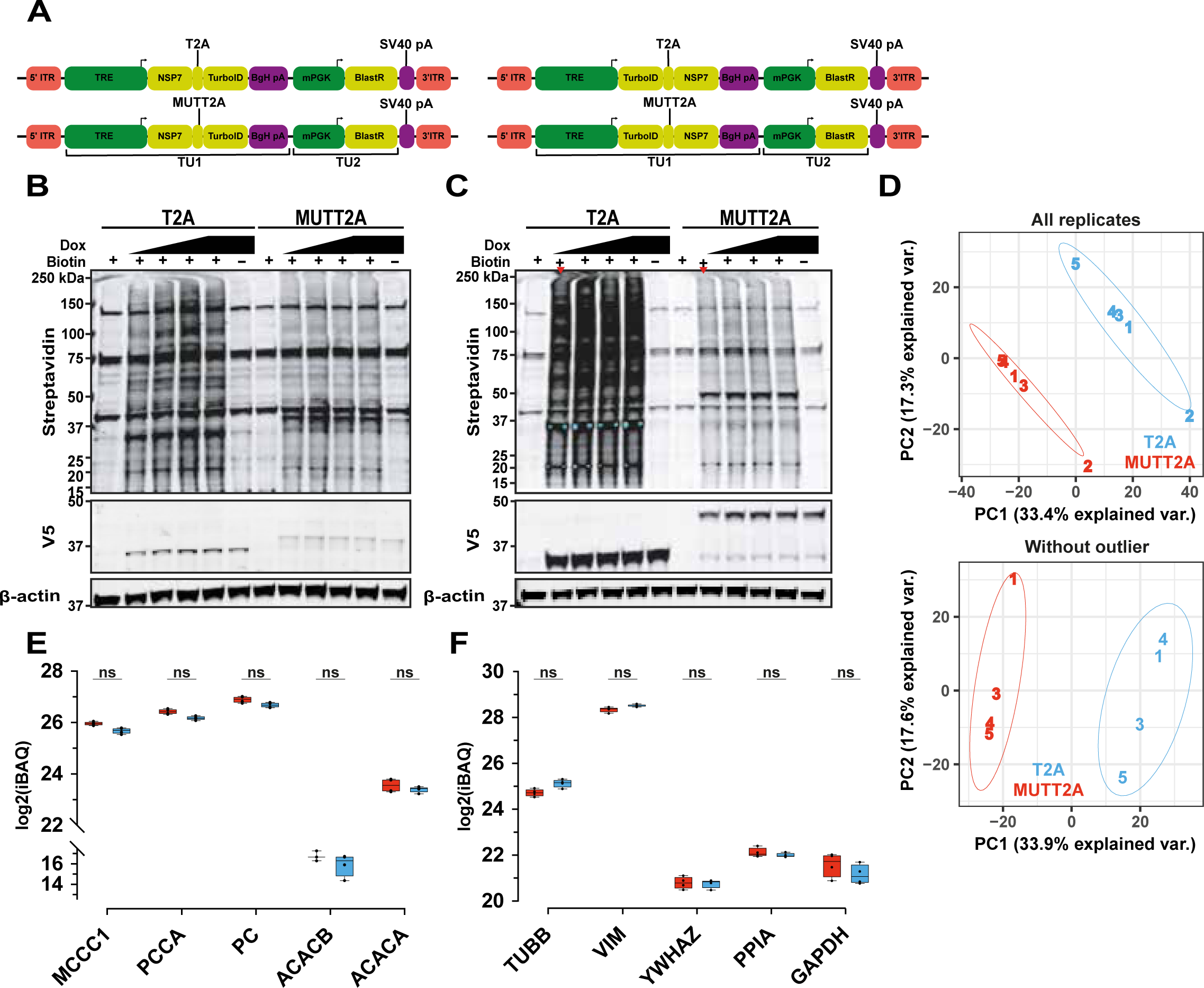
Engineering and validation of TurboID-T2A/MUTT2A-NSP7 A549 cell lines. (a) Schematic overview of N- and C-terminally tagged NSP7 with T2A/MUTT2A-TurboID piggyBac transposon construcs. (b,c) Expression (V5) and biotinylation (streptavidin) staining of A549 cells engineered with constructs shown in (a). (d) Principal component analysis before and after removing outlying replicates. (e) iBAQ intensities of endogenously biotinylated proteins. (f) iBAQ intensities of housekeeping proteins. ITR, inverted terminal repeat; TRE, tetracycline responsive element; GR, glucocorticoid receptor; BgH pA, Bovine growth hormone polyadenylation signal; mPGK, murine phosphoglycerate kinase 1 promoter; BlastR, blasticidine resistance gene; TU, transcriptional unit; Dox, doxycycline; Dex, dexamethasone; PC, principal component; iBAQ, intensity-based absolute quantification.

**Fig. S5.**
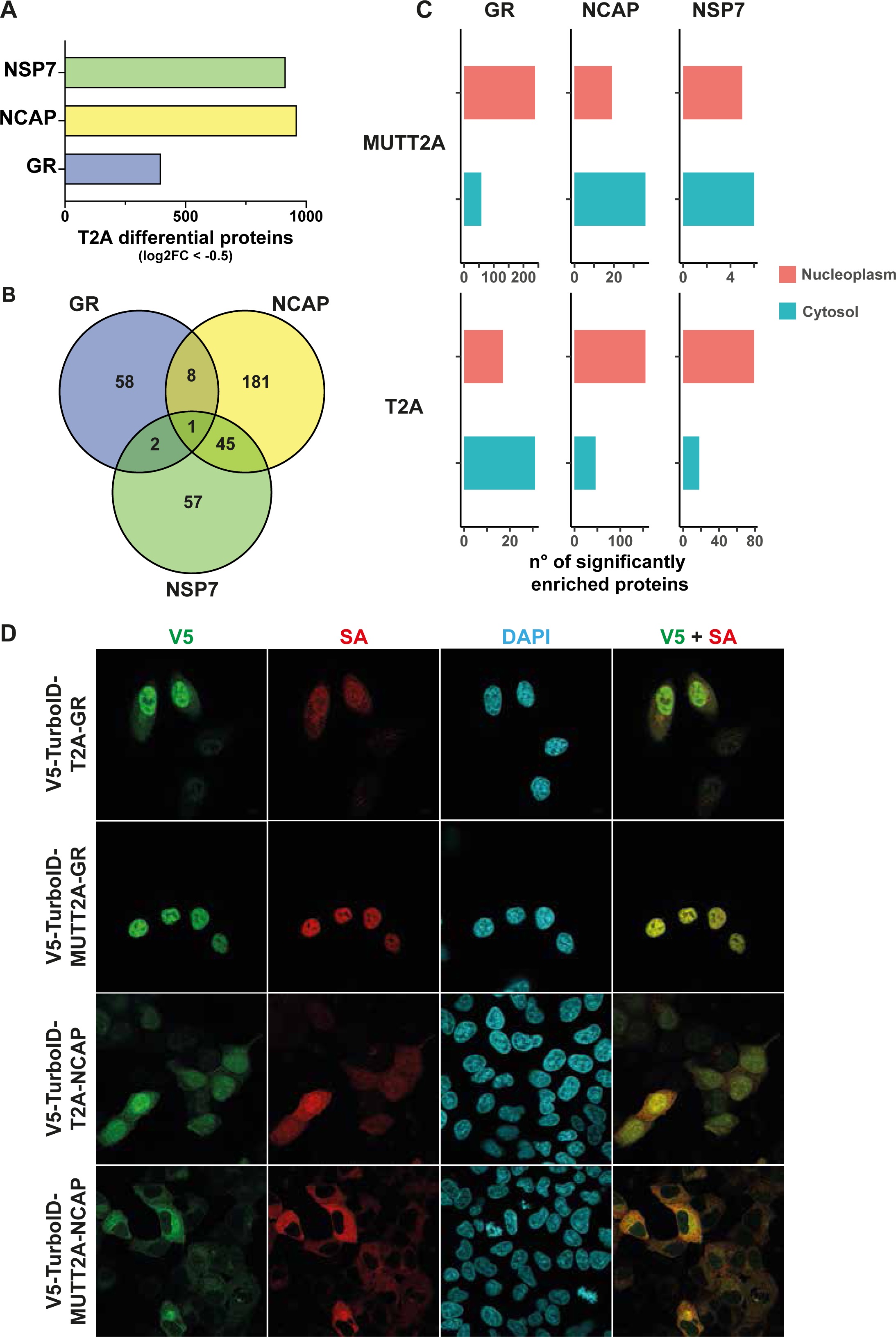
Subcellular localization of TurboID results in T2A skew for cytosolic POIs. (a) Number of differential proteins in the T2A setup (log2FC < −0.5) for NSP7, NCAP, and GR T2A-split link experiments. (b) Overlap of significantly (adj. P-value < 0.05) enriched proteins in the T2A setup between all three experiments. (c) Subcellular localization of all significantly enriched proteins in the T2A and MUTT2A setups. (d) Immunofluorescence of V5-TurboID-T2A/MUTT2A for GR and NCAP. All cell lines were induced with optimized doxycycline conditions and 50 uM biotin. For the GR cell lines, cells were additionally treated with 1 uM dexamethasone. SA, streptavidin; DAPI, 4’,6-diamidino-2-phenylindole.

**Fig. S6.**
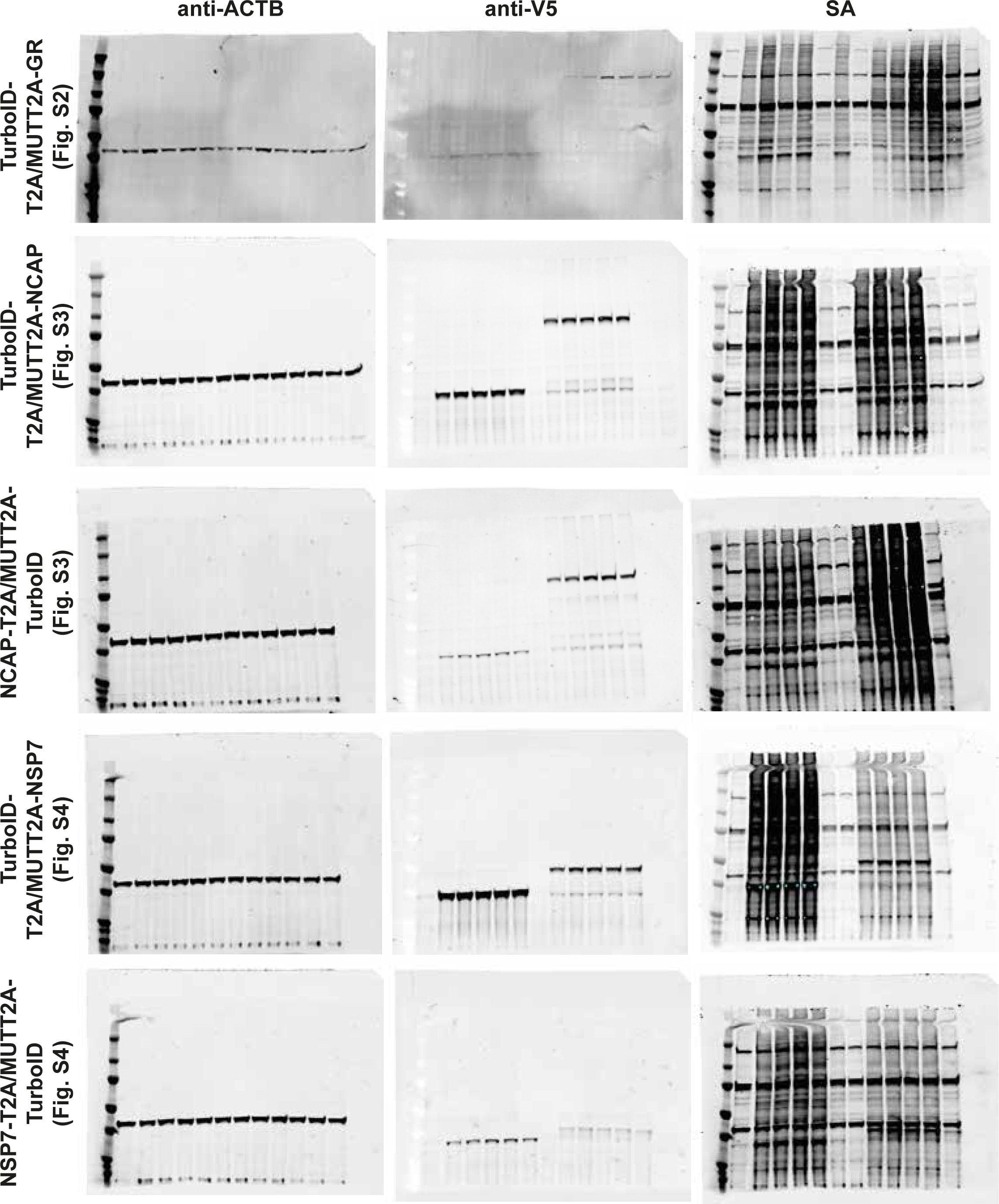
Uncropped immunoblots related to Fig. S2, S3, and S4.

